# Subcellular location defines GPCR signal transduction

**DOI:** 10.1101/2022.12.12.520050

**Authors:** Arthur Radoux-Mergault, Lucie Oberhauser, Simone Aureli, Francesco Luigi Gervasio, Miriam Stoeber

## Abstract

G protein-coupled receptors in intracellular organelles can be activated in response to membrane permeant ligands, which contributes to the diversity and specificity of agonist action. The opioid receptors (ORs) provide a striking example, where opioid drugs activate ORs in the Golgi apparatus within seconds of drug addition. Till date, our knowledge on the signaling of intracellular GPCRs remains incomplete and it is unknown if the downstream effects triggered by ORs in plasma membrane and Golgi apparatus differ. To address this gap, we first assess the recruitment of signal transducers to ORs in both compartments. We find that Golgi-localized ORs couple to Gαi/o probes and are phosphorylated by GPCR kinases (GRK2/3), but unlike plasma membrane receptors, do not recruit β-arrestin or a specific Gα probe. Subsequent molecular dynamics simulations with OR–transducer complexes in model bilayers mimicking plasma membrane or Golgi composition reveal that the lipid environment promotes location selective coupling. Unbiased global analyses then show that OR activation in the plasma membrane and Golgi apparatus has strikingly different downstream effects on transcription and protein phosphorylation. Taken together, the study delineates OR signal transduction with unprecedented spatial resolution and reveals that the subcellular location defines the signaling effect of opioid drugs.

## Introduction

Cells sense changes in their environment through the actions of G protein-coupled receptors (GPCRs) that detect incoming stimuli and transmit the signal by initiating intracellular responses. GPCR-mediated signal transduction plays critical and ubiquitous roles in normal physiology, and aberrations in GPCR pathways can cause disease, making GPCRs key therapeutic targets (*1, 2*). Recent methodological advances, including the use of optical biosensors, have allowed investigating GPCR activation and signaling with unprecedented spatial and temporal resolution on the level of individual cells (*3–5*). This has led to accumulating evidence that some agonists bind and activate GPCRs in intracellular organelles in addition to the plasma membrane (PM), and has established a discrete layer of functional selectivity (*6–8*). An essential step for understanding the diversity of responses triggered by individual ligands is to gain insights into GPCR signaling at different subcellular locations. It remains largely unknown if GPCRs in internal organelles can couple to the canonical PM signal transduction machinery, or alternatively, exhibit a location-selective interaction profile to drive unique downstream responses. Here we start to address these gaps by investigating the compartmentalized signaling of Gi/o-coupled opioid receptors (ORs), which are prototypical members of the rhodopsin-like GPCR family.

ORs are the targets of endogenous neuromodulatory neuropeptides and of therapeutically important analgesic opioid drugs, such as morphine and fentanyl. Nanobody-based sensors recently revealed that mu (MOR), delta (DOR), and kappa (KOR) ORs undergo ligand-dependent activation not only in the PM, but also in different cellular organelles, including endosomes and the Golgi apparatus (*7, 9, 10*). Opioid drugs and opioid neuropeptides strikingly differ in their subcellular activation patterns in that small molecule drugs uniquely trigger activation of ORs in the Golgi apparatus (*7*). Opioid drug access to the cell interior is likely achieved by free diffusion of the small, permeant molecules across cellular membranes. Neurons are known to contain ORs in the PM and in intracellular organelles, including the endoplasmic reticulum and Golgi apparatus, under basal conditions (*11, 12*). Transgenic mice expressing labeled ORs at endogenous levels have revealed varied but often predominant intracellular OR pools across brain regions and neuronal types (*13–16*). Previously, the importance of the internal biosynthetic ORs has mainly been attributed to the ability to replenish PM receptors, however, the recent findings suggest that Golgi-localized ORs may initiate opioid drug-driven responses and have signaling functions.

Elements of the GPCR signaling cascade comprise both spatially confined and diffusing molecules, including transducers, effectors, and second messengers that can act at varying distances relative to the site of ligand-receptor binding. The rapid and dynamic signal propagation makes it difficult to trace back downstream signaling readouts to a precise subcellular site of GPCR activation. Studies that followed rapid receptor-proximal signaling events started to provide location-resolved insights and uncovered Gs-mediated cAMP responses of several GPCRs in endosomes and Golgi apparatus, in addition to the PM (*6, 17–20*). Other studies restricted GPCR activation or signal initiation to a given subcellular site with pharmacological or light-controlled tools, and identified phosphorylation changes, effector activation, and transcriptional changes downstream of intracellular Gs-coupled GPCRs (*19, 21–23*). However, little is known about Gi/o-coupled receptor signaling at internal organelles.

Here we probe MOR and DOR signaling with unprecedented spatial resolution by applying unique pharmacological tools to isolate OR activation in the Golgi apparatus from activation in the PM. First, we demonstrate that canonical GPCR-interacting proteins including G protein-based probes, GPCR kinases (GRKs), and β-arrestin show unique engagement profiles with ORs in the Golgi apparatus, revealing location-selectivity of specific receptor-proximal coupling events. Given that the lipid composition of the PM and Golgi apparatus varies remarkably, we then test if differences in membrane lipids can contribute to the observed coupling selectivity. Using molecular dynamics simulations, we analyze the interactions of two distinct G protein probes with ORs embedded in membranes with a PM- or Golgi-like lipid composition. We find that lipids can directly influence OR coupling and reveal that PM phospholipids interact with a G protein probe to promote a stable OR interaction, while coupling of the same protein is altered in the Golgi-like bilayer. We then measure the downstream signaling effects promoted by PM- or by Golgi-localized OR activation by unbiased global gene expression and protein phosphorylation analyzes and find that unlike OR activation in the PM, Golgi-driven OR activation does not alter gene expression but drives a unique signaling response involving protein phosphorylation.

Taken together, the study identifies location-selective OR signal transduction, which occurs at the receptor-proximal level and results in distinct downstream effects at later time points. Our results further reveal that lipids can directly regulate OR–transducer coupling, providing a possible mechanism to achieve location-selective GPCR responses. Given the highly conserved coupling mechanisms across the GPCR family, the spatial differences that we delineate here for ORs may be widely applicable to signal propagation of GPCR in different cellular locations.

## Results

### Golgi-localized opioid receptors recruit Gαi/o sensors in response to permeant agonists

Activation of ORs residing in the Golgi apparatus has previously been detected using active state-selective nanobodies that bind to Golgi-localized ORs in response to permeant opioid ligands (*7, 9*). We now aimed to test, if the activated internal receptors engage canonical GPCR transducers including G proteins, GPCR kinases, and β-arrestin. First we assessed if Golgi-localized MOR and DOR can interact with G proteins and for this, we employed mini-G (mG) proteins, which are engineered GTPase Ras-like domains of Gα proteins that preserve the molecular contacts formed between the active GPCR and the G protein (**Figure 1A**) (*24, 25*). Since ORs couple to Gi/o, we utilized mG proteins derived from the Gαi/o subunits. For this, we developed three new mG probes based on Gαi1, Gαi2, and Gαi3, which contain deletions and mutations to remove the Gαi membrane anchor, the α-helical domain, and improve protein stability, based on the previously generated mGo probe **(Suppl. Figure S1A)** (*26*). The mGi1, mGi2, mGi3, and mGo proteins were well expressed in HeLa cells, albeit at different levels **(Suppl. Figure S1B)**. We co-expressed mRuby2-tagged mGi1 with GFP-tagged ORs in HeLa cells and tested if the mGi1 probe, which was diffusely distributed in the cytoplasm at basal state, would relocalize to activated receptors in different cellular compartments. mGi1 was recruited to DOR and MOR in the PM after adding peptide agonists (DADLE or DAMGO) or permeant small molecule agonists (SNC80 or fentanyl), as observed using live cell confocal microscopy (**Figure 1B&D)** and total internal reflection fluorescence microscopy (TIR-FM) **(Suppl. Figure S1C&D)**. To quantify mGi1 recruitment to the Golgi pool of ORs, we labeled the Golgi apparatus in living cells with the specific marker ManII-BFP, and quantified mGi1 localization before and after agonist addition. The small molecule agonists SNC80, ARM390, fentanyl, and morphine strongly induced the recruitment of mGi1 to Golgi-localized DOR or MOR (**Figure 1C&E)**, which was dependent on the ligand concentration **(Suppl. Figure S1E)**. Peptide-based agonists did not promote mGi1 recruitment to the Golgi apparatus, confirming the inaccessibility of Golgi-residual ORs to peptide ligands (**Figure 1C&E)** (*7*). We extended our live cell assays to the other Gi/o-based proteins and co-expressed mGi2, mGi3, or mGo with DOR or MOR. Similar to mGi1, the probes had a cytosolic localization in unstimulated cells, and were recruited to PM- and Golgi-localized ORs selectively after treatment with permeant agonists **(Suppl. Fig. S2A-C).**

**Figure 1:**
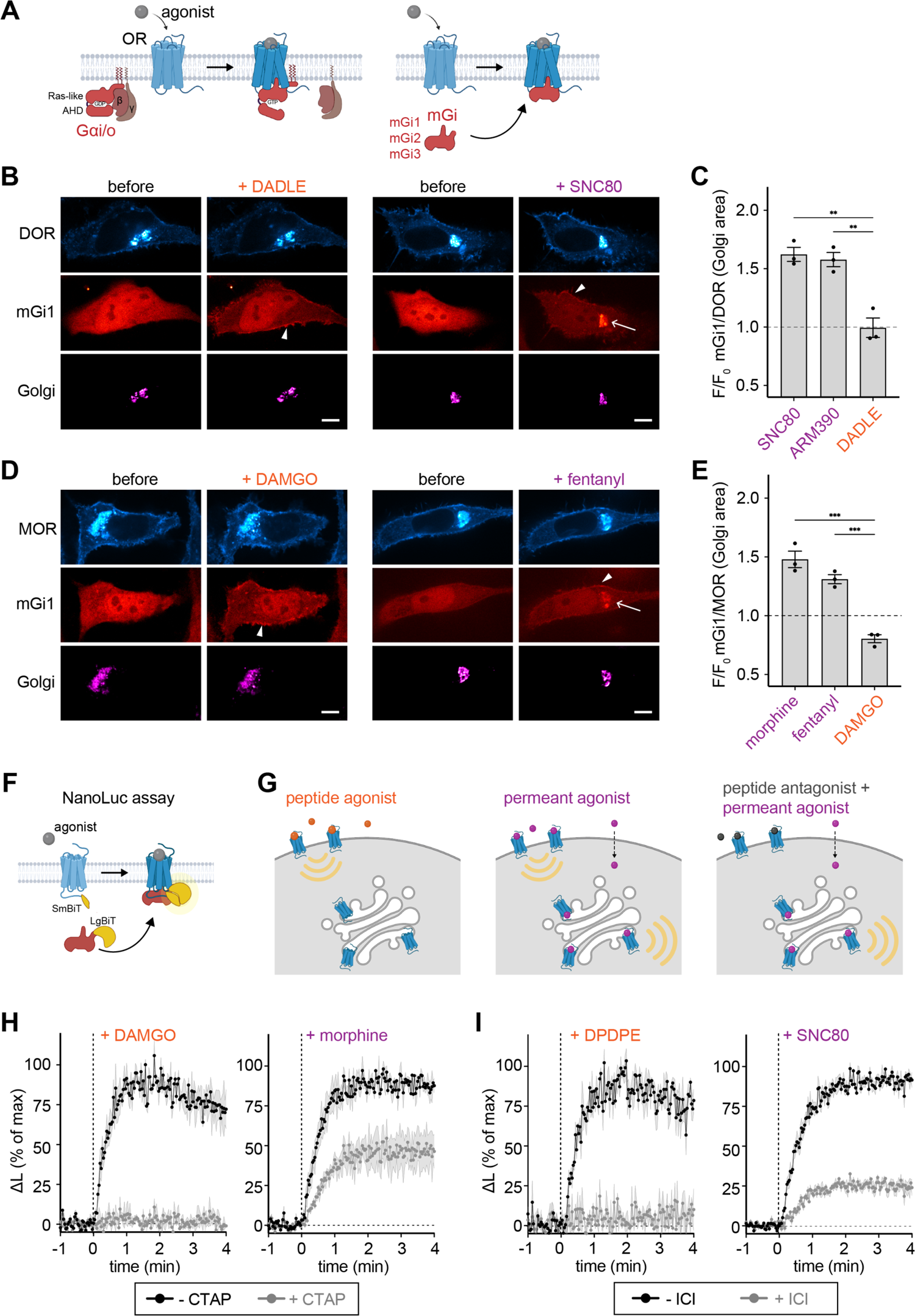
Novel Gαi probes bind ligand activated ORs in the Golgi apparatus. **(A)** Schematic of OR coupling to heterotrimeric Gi/o proteins (left) and to mGi1,2,3 proteins generated in this study (right). Similar to other miniG proteins, the alpha-helical domain (AHD) of Gα is deleted and the Ras-like domain modified, resulting in cytosolic probes that are recruited to the site of OR activation. See also Fig. S1. **(B)** Confocal images of living HeLa cells expressing DOR-SEP (cyan), mGi1-mRuby2 (red), and the Golgi marker ManII-BFP (magenta) before and 5 min after 10 µM DADLE or SNC80 addition. Scale bar = 10 μm. See also Fig. S1 & S2. Arrowheads depict mGi1 at PM, arrow depicts mGi1 in Golgi area. **(C)** Quantification of mGi1-mRuby2 recruitment to DOR in Golgi area (ManII-labeled) upon agonist addition (normalized to Golgi DOR signal). Ligands used at 10 µM. N=3 with > 20 cells analyzed, mean +/- SD. ***p =0.0002 by ordinary one-way ANOVA. **(D)** Confocal images of living HeLa cells, expressing MOR-GFP (cyan), mGi1-mRuby2 (red), and ManII-BFP (magenta) before and 5 min after 10 μM DAMGO or morphine addition. Scale bar = 10 μm. See also Fig. S2. Arrowheads depict mGi1 at PM, arrow depicts mGi1 in Golgi area. **(E)** Quantification of mGi1-mRuby2 recruitment to MOR in Golgi area (ManII-labeled) upon agonist addition (normalized to Golgi MOR signal), using 10 µM DAMGO or morphine, or 1µM fentanyl. N=3 with > 20 cells analyzed, mean +/- SD. **p =0.001 by ordinary one-way ANOVA. **(F)** NanoLuc assay to measure mGi–ORs interaction in living cells. The OR C-terminus is fused to SmBiT and mGi is fused to LgBiT. mGi–OR interaction drives NanoLuc complementation that produces lumiescence in the presence of furimazine. **(G)** Ligands used to selectively activate ORs in the PM (non-permeant peptide agonists), in PM and Golgi apparatus (permeant small molecule agonists), or only in the Golgi apparatus (combination of excess peptide antagonist and permeant agonist). **(H)** Kinetics of mGi1 recruitment to MOR upon DAMGO (100 nM) or morphine (100 nM) addition, with or without prior incubation with peptide antagonist CTAP (10 µM). Mean ± SEM of luminescence signal normalized to untreated cells and to the signal prior to agonist addition. N=3. See also Fig. S2. **(I)** Kinetics of mGi1 recruitment to DOR upon DPDPE (100 nM) or SNC80 (100 nM) addition, with or without prior incubation with peptide antagonist ICI 174,864 (100 µM). Mean ± SEM, analysis similar to (H). N=3. See also Fig. S2.

To determine the kinetics of mGi/o recruitment to ORs and measure the Golgi-localized component of the response, we employed a split nanoluciferase (NanoLuc)-based complementation assay (**Figure 1F**) (*27*). We fused the intracellular C-terminus of DOR and MOR with the small subunit of NanoLuc (SmBiT, 13 aa) and the mGi probes with the larger subunit (LgBiT, 158 aa). HeLa cells expressing OR-SmBiT and mGi1-LgBiT were incubated with the NanoLuc substrate furimazine, and luminescence kinetics were recorded following addition of different ligands or ligand combinations (**Figure 1G**). Adding the peptide agonists DAMGO and DPDPE promoted robust mGi1 recruitment to activated MOR and DOR, with a rapid onset of t_1/2_ < 30 sec (**Figure 1H&I, left)**. The recruitment signal was abolished when cells were pre-incubated with excess of the peptide antagonists CTAP or ICI 174,864 (ICI) to pharmacologically block PM-localized ORs (**Figure 1H&I, left)**. Adding the small molecule agonists morphine and SNC80 led to engagement of mGi1 with MOR and DOR with a similarly rapid onset (**Figure 1H&I, right)** but the maximal luminescence signal was higher than for peptide agonists **(Sppl. Figure S2D&E)**. When cells were pre-incubated with 1000-fold molar excess of peptide antagonists, we found that a significant fraction of the OR–mGi1 interaction driven by morphine or SNC80 remained (**Figure 1H&I, right)**. The results show that MOR and DOR bind mGi1 at intracellular organelles and that the recruitment occurs independently of receptor activation at the PM. Given that the Golgi apparatus is the main site containing intracellular ORs in our experimental system (**Figure 1B&D, Suppl. Figure S2**), we conclude that Golgi-localized DOR and MOR are G protein coupling competent and contribute a significant fraction of the OR–mGi interaction promoted by permeant small molecule agonists.

### GRK2/3 phosphorylate activated GPCRs in the Golgi apparatus

ORs in the PM are regulated through interaction with and phosphorylation by G protein-coupled receptor kinases (GRKs), which modulate downstream responses as they promote β-arrestin binding and concomitant termination of G protein signaling (*28*). The GRKs 2 and 3 (GRK2/3) primarily mediate agonist-driven OR phosphorylation in serine and threonine residues in the cytoplasmic C-terminal tail (*29–31*). We tested if Golgi-resident ORs are phosphorylated in response to permeant agonists. We used a phosphosite-specific antibody that detects MOR when phosphorylated at serine 375, a primary target site for GRK2/3. In basal conditions, cells showed a low signal of MOR-pS375 in both PM and Golgi apparatus (**Figure 2A**). Adding the permeant ligand morphine drove a significant increase in MOR phosphorylation within minutes of agonist application at the PM and also in the Golgi area labeled with ManII-BFP (**Figure 2A&B)**. Upon agonist washout and antagonist (naloxone) addition, MOR phosphorylation at the Golgi apparatus was reversed within 10 min, concomitant with the reversal of receptor dephosphorylation at the PM (**Figure 2A&B)**. Fentanyl also induced phosphorylation of MOR at both PM and Golgi. The peptide agonist DAMGO did not promote phosphorylation of Golgi-resident MOR, while driving phosphorylation of PM receptors, showing that local activation is required for MOR phosphorylation at the Golgi apparatus (**Figure 2C, Suppl. Figure S3A)**. When cells were pre-incubated with the peptide antagonist CTAP to block PM receptors, morphine still drove pronounced phosphorylation of Golgi-resident MOR, revealing that regulation of MOR via phosphorylation occurs locally and independently of PM signaling (**Figure 2D**). When we pre-treated cells with the GRK2/3 inhibitor Compound 101 (Comp101, 30 uM), no morphine-driven MOR-pS375 signal was detected at the Golgi apparatus, which identified GRK2/3 as the likely kinases that mediate MOR S375 phosphorylation at PM and Golgi (**Figure 2E**).

**Figure 2:**
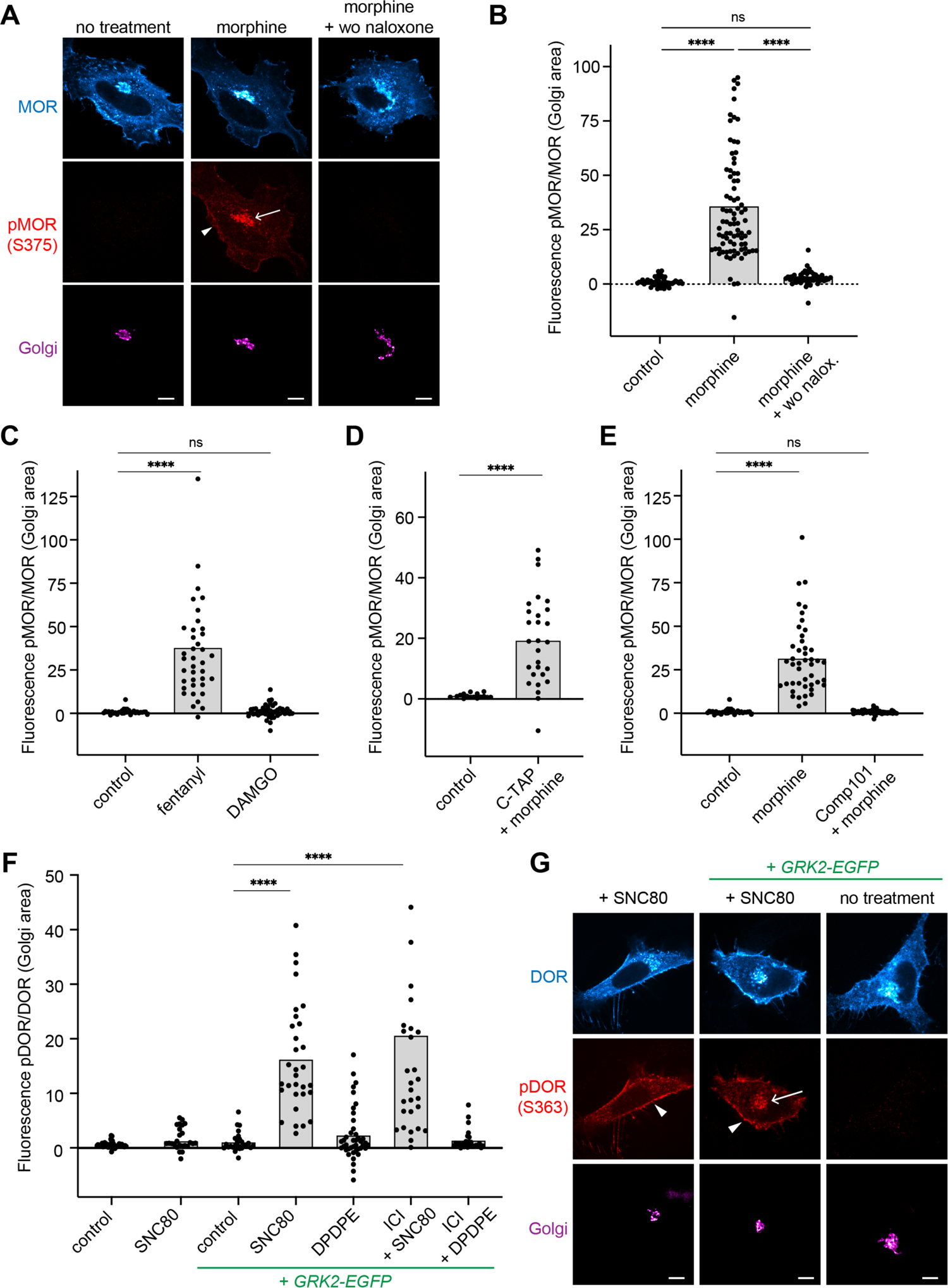
GRK2/3 phosphorylate activated ORs in the Golgi apparatus. **(A)** Confocal images of HeLa cells expressing FLAG-MOR and ManII-BFP (magenta). Cells were fixed, permeabilized and immunolabeled with anti-FLAG (cyan) and anti-pMOR-S375 (red) antibodies. Cells were treated with 10 μM morphine for 5 min. In wash-out condition (wo, right), 10 μM morphine was washed out after 5 min and medium with 100 µM Naloxone added for 10 min. Arrow depicts pMOR in the Golgi area, arrowhead depicts pMOR at the PM. Scale bar = 10 μm. **(B)**, **(C)**, **(D)**, **(E)** Quantification of pMOR/MOR fluorescence in Golgi area (ManII-labeled). Cells were transfected and stained as in (A). Data is normalized to signal of untreated control cells. **(C)** DAMGO used at 10 µM, fentanyl at 1 µM. See also Fig. S3A. **(D)** Cells were pretreated with 10 µM CTAP for 5 min, followed by 100 nM morphine for 5 min. ****p<0.0001 by Welch’s t test. **(E)** 30 µM Comp101 (GRK2/3 inhibitor) was added 45 min prior to adding 10 µM morphine for 5 min. N=3 with cells analyzed >30. ****p<0.0001 by ordinary one-way ANOVA for (B), (C), (E). **(F)** Quantification of pDOR/DOR fluorescence in Golgi area (ManII-labeled). Cells were transfected and stained as in (G). Data normalized to signal of untreated control cells. Cells were treated with 10 μM SNC80 or DADLE. In ICI conditions, 100 μM ICI was added 10 min prior to treatment with 100 nM SNC80 or DADLE. Conditions with GRK2-EGFP co-expression are labeled. N=3 with cells analyzed >25. ****p<0.0001 by ordinary one-way ANOVA. **(G)** Confocal images of HeLa cells expressing FLAG-DOR and ManII-BFP (magenta). Cells were fixed, permeabilized and immunolabeled with anti-FLAG (cyan) and anti-pDOR-S363 (red) antibodies. Treatment with 10 μM SNC80 for 5 min of cells with or without GRK2-GFP expression. Arrow depicts pDOR in the Golgi area, arrowheads depict pDOR at the PM. Scale bar = 10 μm. See also Fig. S3B&C.

We next tested if the Golgi-resident pool of DOR also underlies regulation by GRK2/3 and probed DOR phosphorylation using a phosphosite-specific antibody against serine 363 in the receptor C-tail. We initially did not detect a significant increase in phosphorylation of Golgi-localized DOR after adding the permeant ligand SNC80, while the signal of phosphorylated DOR in the PM strongly increased (**Figure 2F&G)**. We then performed similar immunostainings in cells over-expressing GRK2-GFP. At higher GRK2 levels, a significant phosphorylation of both PM- and Golgi-resident DOR was detected following SNC80 treatment (**Figure 2F&G)**. As for MOR, DOR phosphorylation was independent of PM signaling as pre-incubation with ICI did not block the Golgi-localized DOR phosphorylation (**Figure 2F**). The peptide agonist DPDPE did not drive phosphorylation of DOR in the Golgi apparatus (**Figure 2F**).

Finally, we tested whether the Golgi-pool of the β2-adrenergic receptor (β2AR), a prototypical Gs-coupled GPCR, was also regulated by phosphorylation following their activation. Previous reports indicate that adrenergic receptors in the Golgi apparatus can be activated in HeLa cells upon addition of epinephrine and its derivatives (*6*). Similar to DOR, endogenous kinase levels did not lead to detectable agonist-driven phosphorylation of β2AR (serines 355/356) in the Golgi area, while PM-localized receptors were phosphorylated **(Suppl. Figure S3B&C)**. However, in cells co-expressing GRK2, a pronounced β2AR-pS355/S356 signal was detected for Golgi-resident β2AR minutes after the agonist isoproterenol was applied **(Suppl. Figure S3B&C)**.

The findings show for the first time that the Golgi pools of MOR, DOR, and β2AR are regulated via phosphorylation by GRK2/3 upon activation by permeant ligands. The phosphorylation requires local GPCR activation and occurs independently of PM signaling. In HeLa cells, Golgi-localized, GRK2/3-dependent phosphorylation occurs at endogenous kinase levels for MOR and is detectable at increased GRK2 levels for DOR and β2AR.

### β-arrestin and the mGsi probe show location-selective opioid receptor binding

Since the phosphorylated OR C-tail promotes β-arrestin binding at the PM, we tested if activation of DOR and MOR in the Golgi apparatus recruits β-arrestin2. Using the NanoLuc complementation assay, we tested binding of β-arrestin2 in cells with elevated GRK2 levels to ensure DOR phosphorylation in both the PM and Golgi. As expected, SNC80 treatment drove strong and rapid β-arrestin2 engagement with DOR (**Figure 3A**). However, when PM receptors were blocked with the peptide antagonist ICI-174,864, the β-arrestin2–DOR interaction was abolished (**Figure 3A**), suggesting a lack of β-arrestin binding to Golgi-localized ORs. We next assessed β-arrestin2 recruitment to DOR and MOR using live cell confocal microscopy and found that OR activation with permeant drugs recruited β-arrestin2 to receptors in the PM but not in the Golgi apparatus (**Figure 3B&C, Suppl. Figure S4A)**. We conclude that ligand-activated DOR and MOR in the Golgi apparatus, despite being phosphorylated, do not couple to β-arrestin2.

**Figure 3.**
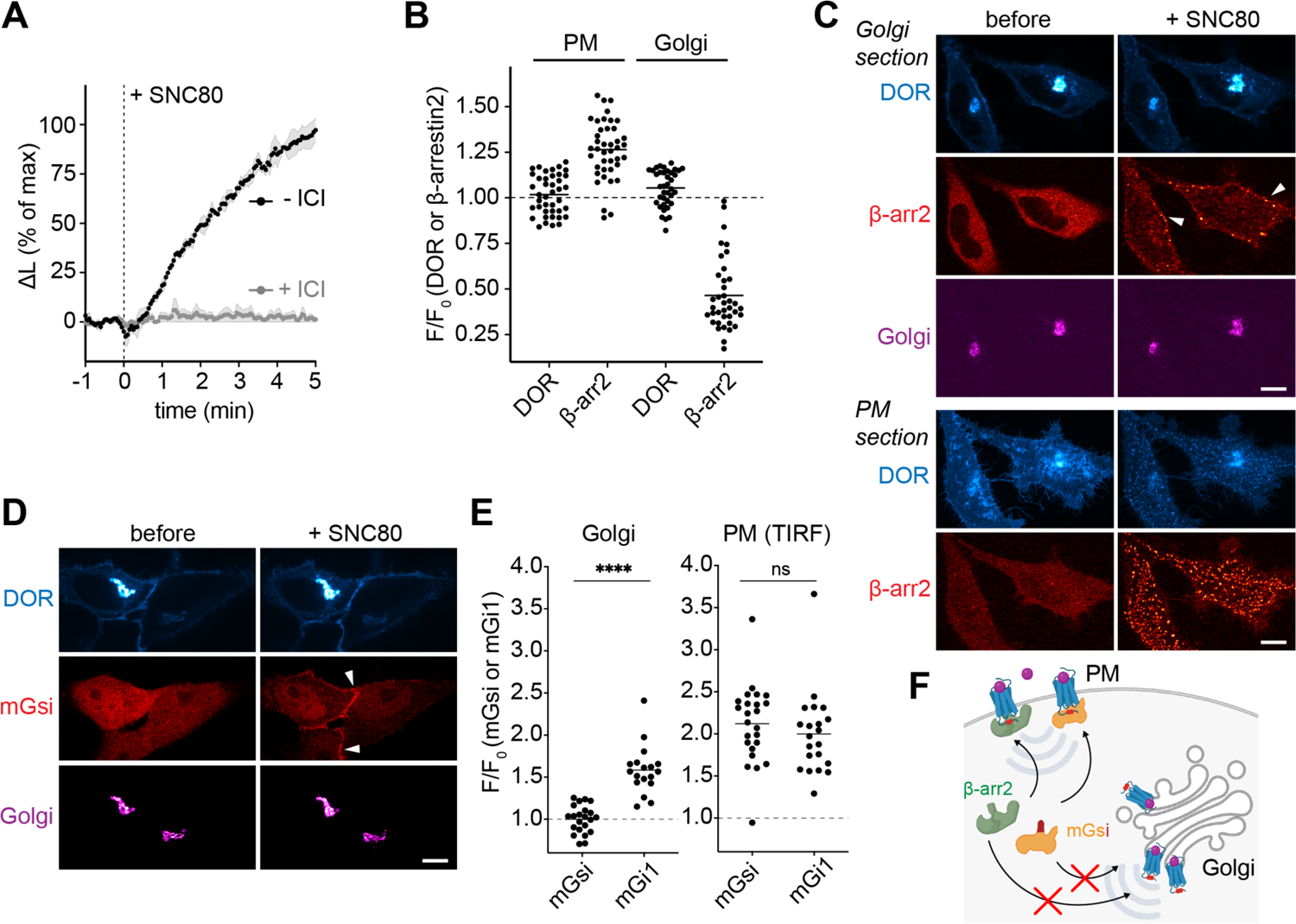
Location-selective OR binding to β-arrestin2 and mGsi. **(A)** Agonist-induced interaction of β-arrestin2-LgBiT with DOR-SmBiT upon adding 100 nM SNC80, in the absence or presence of 100 µM ICI. Change in luminescence signal normalized to untreated cells and to the signal prior to agonist addition. N=3, mean ± SEM. **(B)** Quantification of β-arrestin2-mCherry and DOR signal at the PM or in the Golgi area (ManII-labeled) before and 5 min after 10 µM SNC80 addition. Cells were transfected and images of the same cells acquired with PM or Golgi focus, as described in (C). N=3 with > 35 cells analyzed, mean +/- SD. **(C)** Confocal images of living HeLa cells, expressing SEP-DOR (cyan), β-arrestin2-mCherry (red), and a ManII-BFP (magenta) before and 5 min after 10 µM SNC80 addition. 2 confocal sections of the same cells are shown. Top: focus on the Golgi area, bottom: focus on the PM. Arrowheads depict β-arrestin2 at the PM. Scale bar = 10 μm. See also Fig. S4. **(D)** Confocal images of living HeLa cells, expressing DOR-SEP (cyan), mGsi-mRuby2 (red), and ManII-BFP (magenta) before and 5 min after adding 10 µM SNC80. Arrowheads depict mGsi at the PM. Scale bar = 10 μm. **(E)** Quantification of mGsi-mRuby2 and mGi1-mRuby2 recruitment to DOR in the Golgi or PM. Left: Probe recruitment to DOR in Golgi area (ManII-labeled) as measured by confocal imaging (normalized to Golgi DOR signal). Right: Probe recruitment to the PM as measured with TIR-FM (normalized to DOR signal at PM). Same cells were imaged before and 5 min after adding 10 µM SNC80. N=3 with > 15 cells analyzed, mean +/- SD. ****p<0.0001 by unpaired t test. See also Fig. S4. **(F)** Scheme depicting location-selective recruitment of β-arrestin2 (green) and mGsi (yellow). Both proteins are recruited to ligand activated ORs in the PM but do not bind to active ORs in the Golgi apparatus.

The location-dependent binding of β-arrestin to active ORs was reminiscent of the recruitment pattern of a specific mG probe, termed mGsi, which had recently been shown to bind to active DOR in the PM, but not in the Golgi apparatus (*32*). mGsi consists of the Ras-like domain of Gαs, but contains selectivity determining residues of Gαi in the C-terminal α5-helix **(Suppl. Figure S4B)**, and is widely used as a tool to study Gi/o-coupled GPCR activation and GPCR coupling selectivity (*33, 34*). We hypothesized that the β-arrestin and mGsi engagement with ORs may underlie a shared regulatory mechanism that differs based on the subcellular location, and which is distinct to mGi1 that bound active ORs in both PM and Golgi (**Figure 1**). We also reasoned that comparing OR binding to mGsi and to mGi1, two proteins that have similar overall conformation and sequence, may enable uncovering molecular mechanisms that promote location-selective OR coupling of one protein but not of the other. First, we confirmed the lack of mGsi binding to Golgi-localized DOR upon small molecule agonist (SNC80) addition, while the PM DOR pool engaged mGsi (**Figure 2F&G).** We found the same location-selective engagement pattern when we extended our analyses to MOR. The permeant ligand fentanyl drove mGsi recruitment to active MOR in the PM but not in the Golgi apparatus **(Suppl. Figure S4C)**. To rule out the possibility that the different recruitment patterns of mGsi and mGi1 were caused by differences in the probes’ sensitivities for ORs, we quantified the agonist-driven recruitment of mGsi and mGi1 to cell surface ORs using TIR-FM. Both mGsi and mGi1 were recruited to PM-localized DOR or MOR to the same extent and with similar kinetics (**Figure 3E, Suppl. Figure S4C&D)**. The mGsi-specific location bias suggested that the determinants for OR–G protein coupling in the Golgi apparatus comprise structural features outside the Gαi/o C-terminus, which is shared between the probes. Furthermore, the similar subcellular recruitment pattern of β-arrestin2 and mGsi (**Figure 3F**) suggested a common mechanism, and motivated us to mechanistically explore the location-selective GPCR coupling behavior of the mGsi probe.

### Subcellular membrane composition affects opioid receptor coupling

One clear distinction between the PM and the Golgi apparatus is the composition of lipids that make up the membrane and therefore we set out to test if differences in the local lipid environments contribute to the distinct recognition of ORs by mGsi and mGi1. We used molecular dynamics (MD) simulations to perform a comprehensive molecular assessment of the MOR–mG interactions in model membrane bilayers with PM- or Golgi-like lipid composition and properties (**Figure 4A**). We focused on MOR because the high-resolution cryoEM insights into MOR–Gi coupling provided a valuable basis for our analyses (*35*). The membrane models were built based on previous computational studies of realistic membranes, with multiple lipid types and a high degree of compositional complexity (*36–42*) (**Figure 4A**). The initial structures of the MOR–mGsi and MOR–mGi1 complexes were generated using “*Alphafold2-multimer*” and each complex was embedded in either membrane model by employing CHARMM-GUI (see methods section). All four final structures underwent 1.0 μs long classical MD calculations.

**Figure 4.**
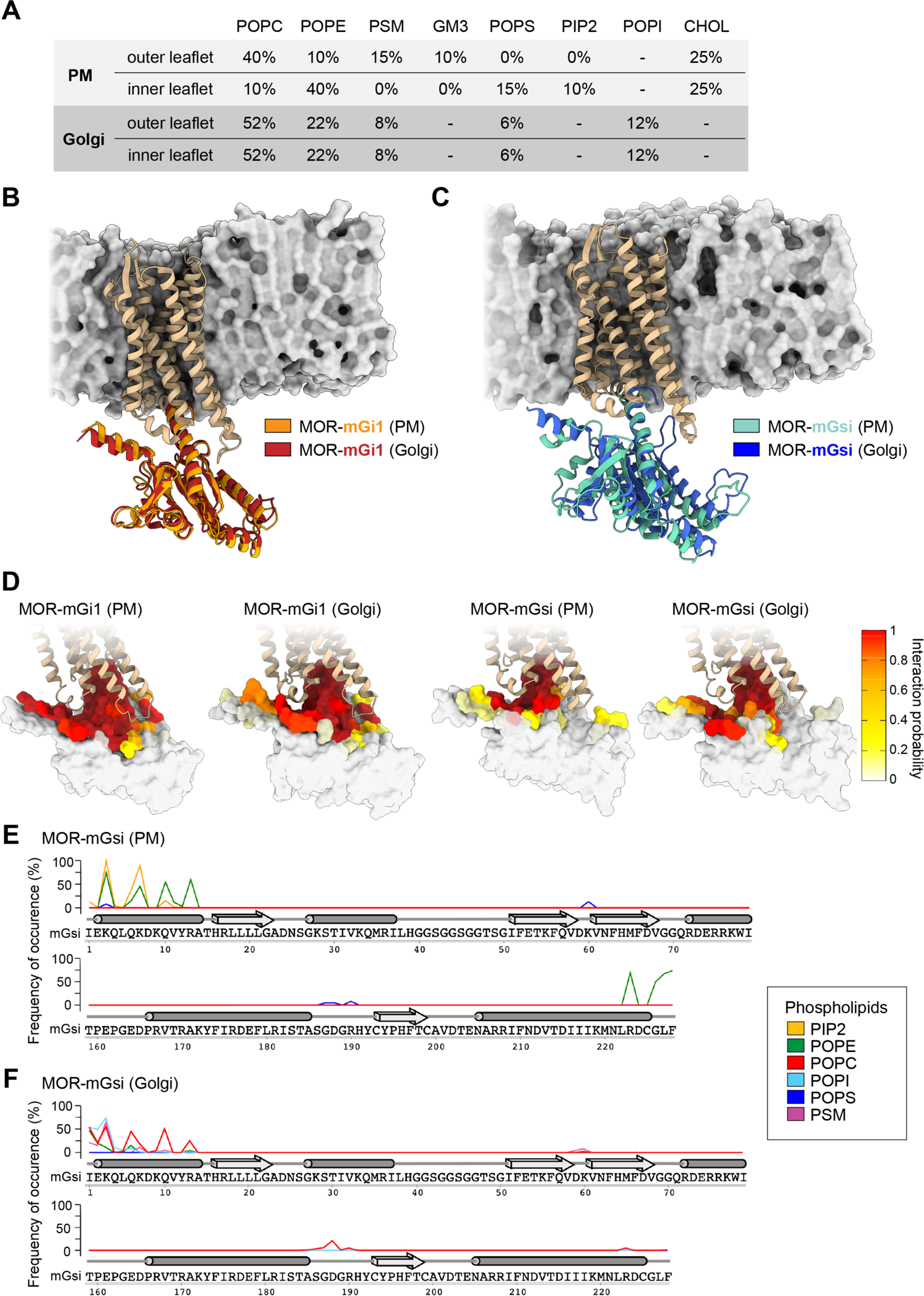
The lipid environment regulates OR – mGsi interaction. Results of the MD simulations carried out with MOR–mGi1 and MOR–mGsi heterodimers in PM and Golgi-like membrane models. **(A)** Lipid composition of the PM and Golgi-like membrane models employed in the MD simulations. 1-palmitoyl-2-oleoyl-sn-glycero-3-phosphocholine (POPC), 1-palmitoyl-2-oleoyl-sn-glycero-3-phosphoethanolamine (POPE), palmitoyl sphingomyelin (PSM), monosialo dihexosyl ganglioside (GM3), 1-palmitoyl-2-oleoyl-sn-glycero-3-phosphoserine (POPS), 1,2-diacyl-sn-glycero-3-phospho-1-D-myo-inositol 4,5-bisphosphate (PIP2), 1-palmitoyl-2-oleoyl-sn-glycero-3-phosphoinositol (POPI), cholesterol (CHOL). **(B)**,**(C)** Superimposition of the most relevant conformations assumed by the MOR– mGi1 and MOR–mGsi heterodimers in the PM and Golgi membranes. MOR is colored in tan. See also Fig. S5. **(B)** mGi1 in the PM environment is orange, mGi1 in the Golgi environment is red. **(C)** mGsi in the PM environment is teal, mGsi in the Golgi environment is blue. Lipid bilayer schematics depicted in the panels are based on the Golgi membrane models, which are displayed through their solvent exposed surface (gray color). **(D)** Frequency of occurrence of the contacts between MOR with mGi1 and MOR with mGsi. mGi1 and mGsi are represented through their solvent exposed surfaces and colored according to the local interaction probability values following the color gradient on the right. See also Fig. S6 and S7. **(E)**,**(F)** Frequencies of occurrence of the contacts between mGsi and phospholipids in the PM **(E)**, and the Golgi membrane **(F)**. The α-helices (tubes) and β-sheets (arrows) of mGsi are displayed on top of the primary sequence. The frequency of interaction of each phospholipid species is colored according to the legend. mGsi residues 70-159 are omitted since no lipid interactions were detected.

First, we assessed the overall dynamics and conformational stability of the OR–mG complexes. For this, we computed the Root Mean Square Deviation (RMSD) of the secondary structure Cα atoms and plotted the RMSD values for both MOR or the mGi1 / mGsi probes as a function of the simulation time **(Suppl. Figure S5A&B)**. MOR exhibited good conformational stability in all four simulations (MOR bound to mGi1 or mGsi, in PM or Golgi membrane) (average RMSD ∼1.0 Å) **(Suppl. Figure S5A)**, however, the mG probes exhibited a more intricate behavior **(Suppl. Figure S5B)**. While mGi1 displayed limited conformational flexibility in both membrane environments (average RMSD ∼1.0 Å), mGsi engaged a stable binding pose only when MOR was embedded in the PM bilayer. In the Golgi-like environment, a higher average RMSD value for mGsi (∼2.2 Å) was observed, suggesting a less favorable MOR– mGsi interaction. A membrane-dependent behavior of MOR–mGsi binding was also detected by cluster analyses of the conformational states visited by the protein complexes during the different MD simulations. We found that MOR–mGi1 heterodimers assumed a similar binding mode in both PM- and Golgi-like membranes (**Figure 4B**), while the MOR–mGsi heterodimers exhibited significantly different binding geometries in the two membranes (**Figure 4C**).

We then investigated whether distinct interaction networks between the receptor and the mG probes might contribute to the different coupling behaviors of mGi1 and mGsi. We computed the frequency of occurrence of the contacts formed between MOR and mGi1 or MOR and mGsi during the simulations. An overall higher degree of engagement between MOR and mGi1 in comparison to mGsi was observed in both membrane models (**Figure 4D**). MOR and mGi1 interacted via an extensive network of salt-bridges **(Suppl. Figure S6**), mediated by mGi1 residues in the C-terminal helix (H5), the N-terminal helix (HN), and the hns1, s2s3, and h4s6 linkers (domains according to GPCRdb), closely paralleling the binding of full length Gαi to MOR (*35*). The MOR–mGi1 interactions were virtually identical in both membrane environments **(Suppl. Figure S6**). In particular, the three salt bridges *mGi1-K171–MOR-E272*, *mGi1-E175–MOR-K273* and *mGi1-D198–MOR-K273* were able to strongly engage helix 6 in MOR and ease the retention of MOR’s active state. In contrast, MOR–mGsi binding was dominated by hydrophobic interactions and characterized by a lower number of salt-bridges that differed according to the membrane environment (**Figure 4D, Suppl. Figure S7**). Beyond the interactions mediated by the mGsi C-terminal helix, the MOR–mGsi complex was held together by three inter-protein salt bridges in the PM environment (*mGsi-R13–MOR-D179*, *mGsi-D157–MOR-K271*, and *mGsi-D215– MOR-K273*), yet only one salt-bridge was detected in the Golgi-like membrane (*mGsi-D215–MOR-K273*) **(Suppl. Figure S7**). The analyses suggest that MOR binds mGsi with lower affinity than mGi1 due to a reduced interaction network, and are consistent with membrane-dependent binding of mGsi to MOR.

To assess if the membrane composition might impact mGsi–MOR binding, we analyzed the frequency of occurrence of contacts between the mGsi residues and each phospholipid species in the PM- and the Golgi-like bilayers. We found that both the N-terminal and C-terminal helices of mGsi bound to PM lipids, yet different lipid species were involved (**Figure 4E**). While both helices engaged with POPE (1-palmitoyl-2-oleoyl-sn-glycero-3-phosphoethanolamine), the positively charged residues in the N-terminal helix specifically bound to PIP2 (phosphatidylinositol-4,5-biphosphate), a highly negatively charged phospholipid that is enriched in the PM. The interactions of mGsi with lipids in the Golgi-like membrane exhibited a different pattern (**Figure 4F**). The C-terminal helix of mGsi interacted poorly with lipids and the N-terminal helix also showed a lower level of membrane engagement, mainly via the zwitterionic phospholipid POPC (1-palmitoyl-2-oleoyl-sn-glycero-3-phosphocholine). The findings identify interactions between mGsi and individual phospholipids that may promote MOR–mGsi complex assembly in the PM, but not in the Golgi environment.

Together, the results suggest a lipid-based mechanism for location-selective MOR coupling and offers a molecular explanation for the lack of mGsi binding to Golgi-localized ORs, which is consistent with the subcellular binding pattern we find in living cells (**Figure 3**). We conclude that OR coupling to specific proteins may require the interplay between receptor-mediated and phospholipid-mediated contacts in complex assembly.

### Opioid receptor signaling at the PM, but not at the Golgi apparatus, alters gene expression

As our results have revealed differences in the molecular interactions between ORs and signaling partners at the PM and the Golgi apparatus, we set out to test if signals initiated in the different cellular compartments result in distinct downstream effects. GPCRs exert their functions by engaging transducers and effectors that produce second messengers and cause phosphorylation changes, which can give rise to changes in gene expression. We profiled OR-mediated signaling using two parallel approaches, transcriptomics and phosphoproteomics. First, we performed differential gene expression analysis using RNA sequencing (RNA-seq) and compared the transcriptome of control cells and cells treated with agonists for 1.5 h or 6h. The signaling studies were performed using HEK293 cells stably expressing DOR (HEK293-DOR), which contained pronounced DOR pools in both the PM and the Golgi apparatus at steady-state (**Figure 5A**). To probe if acute signaling of PM-localized ORs drives robust changes in gene expression, we treated cells with the peptide agonist DPDPE. At 1.5h and 6h, significant transcriptional changes were detected, with differential expression of 34 genes at 1.5 h and 272 genes at 6 h (adjusted p-value < 0.05, fold-change > 1.5 or < −1.5) (**Figure 5B&C, Suppl. Figure 8A&B, Suppl. Table 1**). Consistent with previous reports on transcriptional regulation by ORs (*43–45*), the 1.5 h hits comprised immediate-early response genes (IER3, EGR1, and EGR3), regulators of mitogenic signaling (DUSPs), Hippo pathway genes (CTGF, ANKRD1, CYR61), and transcription factors (e.g. MAFF, FOSL1), and we also detected their upregulation in separate experiments using RT-qPCR **(Suppl. Figure S8C)**. The transcriptional response driven by DOR was distinct to the changes promoted by isoproterenol-activated Gs-coupled β2AR **(Suppl. Figure S8A)**, showing that gene expression changes can reveal specific downstream signals. We then probed whether activated Golgi-localized ORs initiate transcriptional changes. To restrict DOR signaling to the Golgi area, we again used the peptide antagonist ICI at high concentrations to block PM receptors, and subsequently stimulated Golgi-resident ORs by adding the permeant agonist SNC80 for 1.5 h or 6 h (ICI-SNC80 condition). First, we confirmed that ICI treatment alone did not affect cellular gene expression **(Suppl. Figure S8D)**. We then performed live cell confocal imaging of HEK293-DOR cells expressing the OR-activity sensor EGFP-Nb33, which confirmed that the ICI-SNC80 treatment drove Golgi-localized DOR activation that lasted for 6 h, while no DOR activation at the PM was detected (**Figure 5D**). Strikingly, RNA-seq-based differential gene expression analysis identified a complete lack of transcriptional changes upon Golgi-restricted DOR activation at 1.5 h or 6 h (fold-change > 1.5 or < −1.5, adjusted p-value < 0.05) (**Figure 5E&F)**. This was not due to agonist-specific differences, because when we performed RNA-seq analysis of cells treated with SNC80 in the absence of ICI, we detected a pronounced transcriptional response that was identical to the DPDPE effects **(Suppl. Figure S8A&E, Suppl. Table 1**).

**Figure 5:**
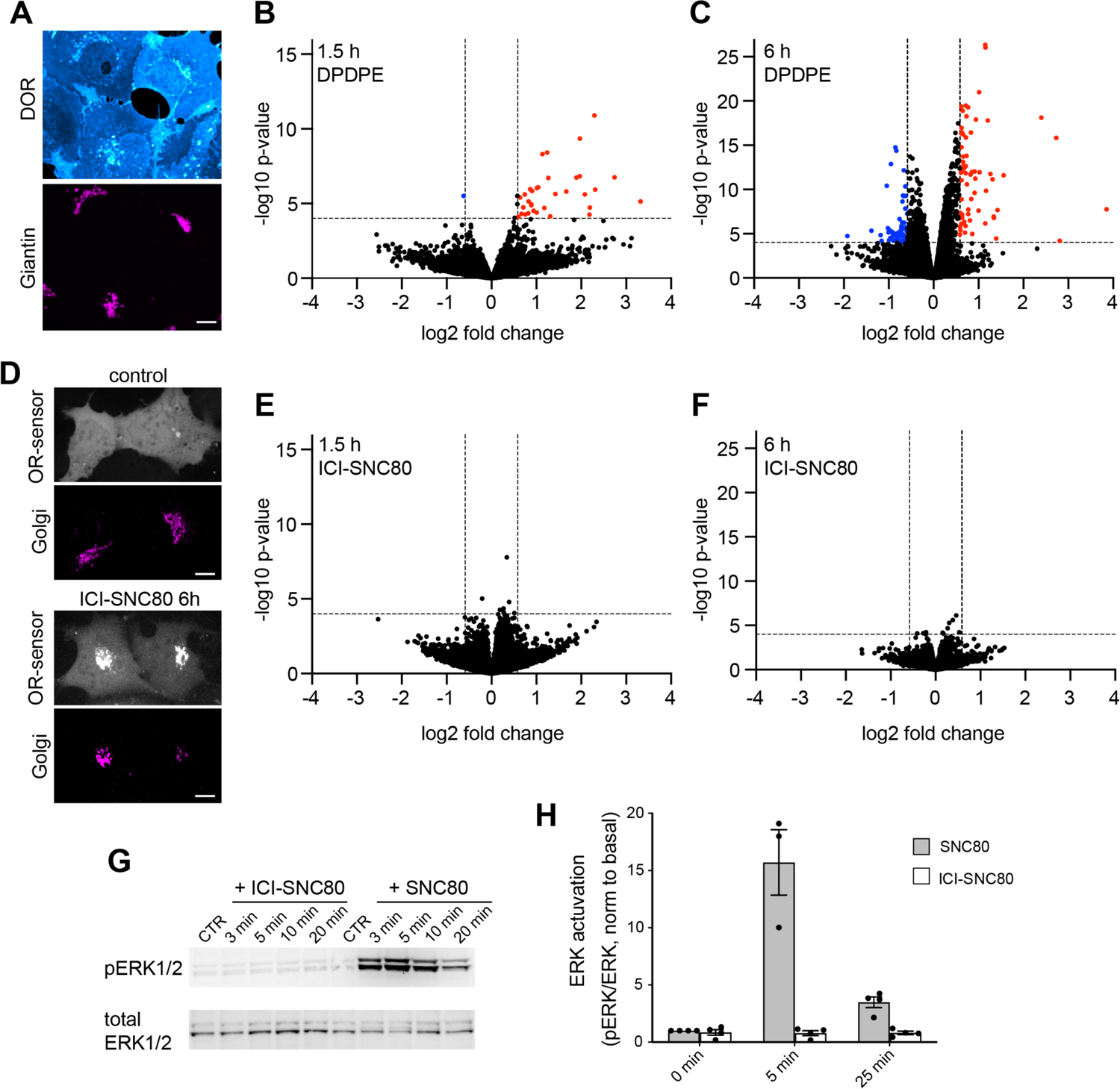
Golgi-restricted DOR activation does not promote a transcriptional response. **(A)** Confocal images of HEK293 cells stably expressing FLAG-DOR (HEK293-DOR). Cells were fixed, permeabilized, and immunolabeled with anti-FLAG (cyan) and anti-Giantin antibodies (magenta). Scale bar = 10 µm. **(B)**,**(C)** Volcano plots of differentially expressed genes in HEK293-DOR cells treated with DPDPE (100 nM) for 1.5 h (B) or 6 h (C) versus mock-treated control cells. Results are presented as the mean differential gene expression from 3 replicates and expressed as log2 fold-change. Genes significantly upregulated are shown in red (QL F-test, FC > 1.5, adj. p-value < 0.05). Genes significantly downregulated are shown in blue (QL F-test, FC < −1.5, adj. p-value < 0.05). See also Fig. S8. **(D)** Confocal images of living HEK293-DOR cells, expressing the OR-activity sensor EGFP-Nb33 (gray) and GalT-DsRed. Depicted are control cells and cells treated for 6 h with ICI (100 µM, 5 min pre-incubation) and SNC80 (100 nM). Scale bar = 10 µm. **(E)**,**(F)** Volcano plots of differentially expressed genes in HEK293-DOR cells treated with ICI-SNC80 (ICI = 100 µM, SNC80 = 100 nM) for 1.5 h (E) or 6 h (F) versus control cells (100 µM ICI treated). Results are presented as the mean differential gene expression from 3 replicates and expressed as log2 fold-change. No genes are significantly changed in their expression (criteria as in B and C). See also Fig. S8. **(G)** Representative immunoblot of ERK1/2 activation upon treatment of HEK293-DOR cells with ICI-SNC80 (100 µM & 100nM) or SNC80 (100nM) for indicated times. **(H)** Quantification of pERK1/2 protein levels normalized to total ERK1/2 in HEK293-DOR cells treated as in (G), for 5 min and 25 min. Mean +/- SEM, N=3.

It is well established that GPCRs, including ORs, regulate expression of primary and secondary response genes through activation of mitogen-activated protein kinase (MAPK) modules. Our RNA-seq results suggested that Golgi-localized ORs may not drive MAPK signaling. We probed ERK1/2 MAPK activity after Golgi-restricted DOR activation (ICI-SNC80) and indeed detected no ERK1/2 phosphorylation (**Figure 5G&H)**. Yet SNC80 treatment in the absence of ICI showed increased ERK1/2 activity downstream of OR activation (**Figure 5G&H)**, showing that transcription regulators are differentially affected by activation of ORs in the PM and Golgi apparatus.

### Activation of opioid receptors in the Golgi apparatus produces unique effects on protein phosphorylation

We next turned to mass-spectrometry-based analyses of the phosphoproteome as a parallel unbiased and global method for profiling cellular signaling and focused on early time points, since GPCR-driven phosphorylation events occur rapidly (*46–49*). Using Liquid Chromatography-Electrospray Ionization-Tandem Mass Spectrometric (LC/ESI-MS/MS) analyses, we captured OR-mediated phosphorylation changes in HEK293-DOR cells treated with SNC80 (PM signaling control) or ICI-SNC80 (Golgi-restricted signaling) for 5 min or 25 min. OR-mediated changes of phosphopeptides were considered significant if a fold change > 1.2 or < −1.2 and p-value < 0.05 was reached, and the corresponding protein level was unchanged. The cutoff was based on the detected fold change of a DOR C-terminal peptide, known to be phosphorylated upon OR activation (site: DOR-p363, 1.26 fold change) (**Figure 6A&B)**. SNC80 treatment induced phosphorylation of 149 and 142 peptides after 5 min and 25 min respectively, with 48 shared phosphopeptides between the time points (**Figure 6A,B,E, Suppl. Table 2**). Golgi-restricted DOR activation by ICI-SNC80 induced phosphorylation of 35 and 82 peptides after 5 min and 25 min treatment respectively, with 13 shared hits between the time points (**Figure 6C,D,E, Suppl. Table 2**). Signaling from the PM and from the Golgi apparatus resulted in largely different phosphorylation effects as only 3 phosphopeptides were shared between the SNC80 and ICI-SNC80 hits (**Figure 6E, Suppl. Table 2**), which is consistent with a unique Golgi-localized OR response.

**Figure 6:**
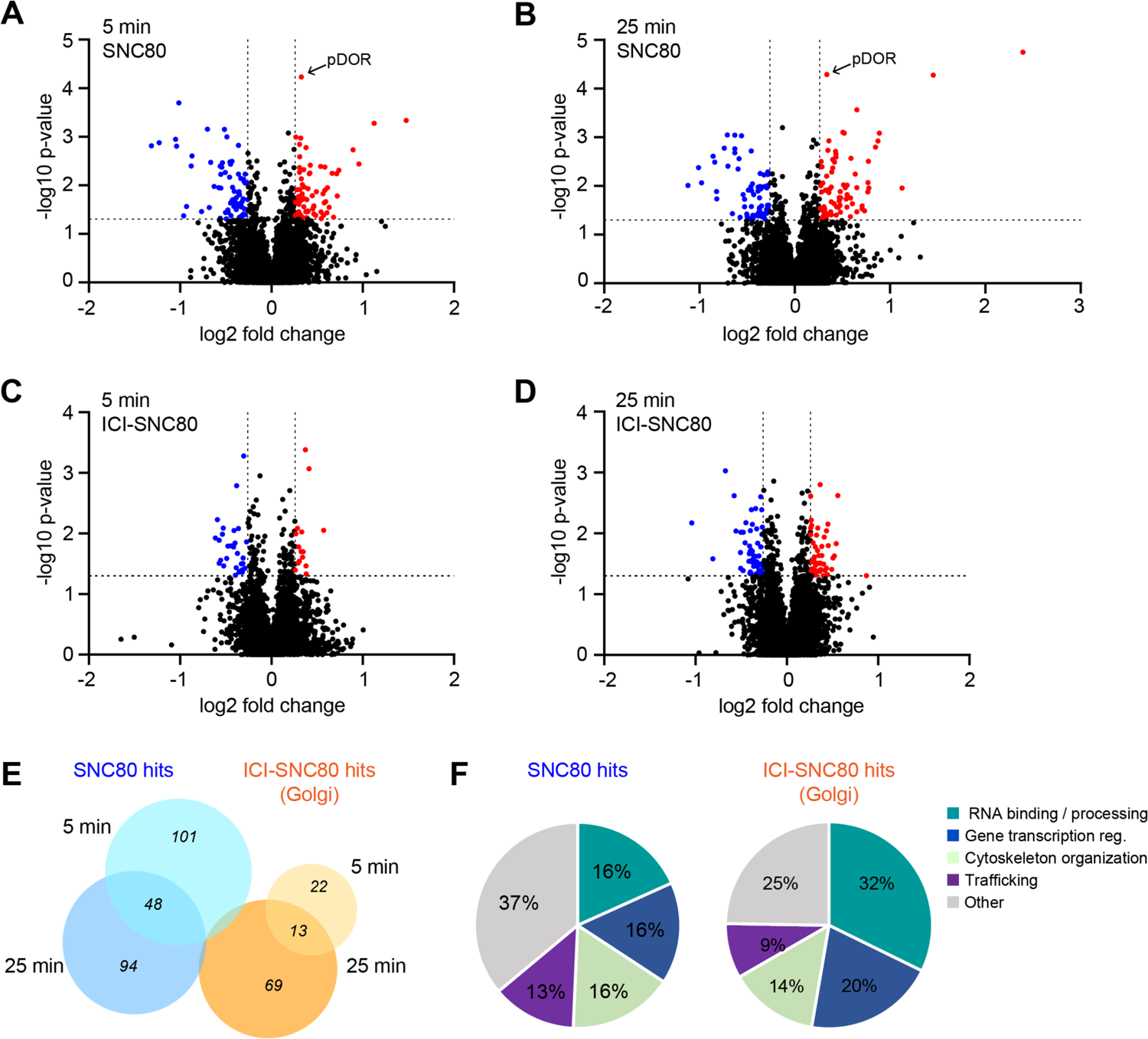
DOR activation in the Golgi apparatus promotes unique changes in the phosphoproteome. **(A)-(D)** Volcano plots of phosphopeptides identified by mass-spec in HEK293-DOR cells treated with SNC80 (100 nM) for 5 min **(A)** and 25 min **(B)** or with ICI-SNC80 (ICI = 100 μM, SNC80 = 100 nM) for 5 min **(C)** and 25 min **(D)** relative to control cells. Results are presented as the mean phosphopeptide abundance ratios between treated and mock-treated cells from 3 replicates and expressed as log2 fold-change. Peptides showing a significantly higher phosphorylation upon treatment are shown in red (ANOVA, FC > 1.2, p-value < 0.05) and peptides with significantly lower phosphorylation are shown in blue (ANOVA, FC < −1.2, p-value < 0.05). **(E)** Venn diagrams showing the overlap between phosphopeptides regulated upon 5 min and 25 min SNC80 treatment or 5 min and 25 min ICI-SNC80 treatment. Only three phosphopeptides are shared between the SNC80 and ICI-SNC80 hits (VIM, ATP2B1, SRRM2). **(F)** Classification of the modulated phosphoproteins downstream of SNC80 or ICI-SNC80 treatment into functional groups based on Metacore (Clarivate), STRING (v. 11.5) and Pubmed and Uniprot curation. See also Fig. S9.

Analyzing the phospho-responses revealed that 25% of regulated proteins downstream of SNC80 treatment have been previously linked to OR function (**Suppl. Table 2**), which provided a validation of the phosphoproteomic dataset. Notably, the mTOR pathway previously detected upon KOR activation in mice brain (*48, 50*), was also affected downstream of DOR activation by SNC80 (SLC3A2, PRKAA2, MTOR, RPS6KA1, BRAF, according to KEGG pathway). In addition, PAK1 and PAK2 were significantly phosphorylated upon SNC80, and previously also identified upon KOR activation in HEK293 cells (*51*). None of these phosphoproteins were, however, regulated downstream of Golgi-restricted OR activation. Similarly, several endocytic proteins were detected upon SNC80 treatment (PARD3, IGF2R, DNM1, EPS15L1, RAB11FIP1, CBL, SNX1, according to KEGG pathway), yet they did not change upon ICI-SNC80 addition. While the ERK1/2 kinases did not pass the cutoff criteria used to define hits, an increase in pERK2 (MAPK1) was detected in the SNC80 condition (t=25 min, FC = 1.18). pERK was not detected after adding ICI-SNC80, consistent with the location-dependent OR effects on ERK1/2 activity described earlier (**Figure 5G&H)**. We then aimed to identify upstream kinases by applying kinase-substrate enrichment analyses (KSEA App), pooling all unique phosphosites regulated by either SNC80 or ICI-SNC80 treatment **(Suppl. Figure S9A-C, Suppl. Table 2**). In the SNC80 condition, ERK1 and ERK2 were predicted as active kinases with highest confidence, and the cAMP-regulated PKA was predicted to be significantly less active **(Suppl. Figure S9A&C, Suppl. Table 2**), pinpointing kinases with known roles downstream of OR signaling. No significant kinase activity could be assigned based on the phosphosites identified in the ICI-SNC80 conditions **(Suppl. Fig. S9B)**. We noted that 5 min after Golgi-localized ORs activation, 75% of regulated proteins showed a reduction in their phosphorylation status, which suggests that internal OR activation may predominantly downregulate a basal kinase tone.

Next, we categorized the phosphoproteins regulated by SNC80 or by ICI-SNC80 treatment according to cellular function (**Figure 6F**). Downstream of Golgi-localized OR activation, 32% of hits had roles in RNA binding and processing, and GO term analysis identified significant enrichment of molecular functions related to RNA regulation in the ICI-SNC80 response (**Suppl. Table 2**). In addition, 20% ICI-SNC80 hits were linked to transcriptional regulation, and several regulated proteins showed functions in cytoskeleton remodeling and membrane trafficking, some of them with established cellular localization at the Golgi apparatus (SNAPIN, ARHGAP21, ALG2, CAMSAP1/3) (**Figure 6F).** The identified hits present novel candidate proteins that may specifically act downstream of the drug-induced Golgi-localized OR activation wave (**Suppl. Table 2**).

Taken together, the unbiased downstream signaling analyses demonstrate that Golgi-localized ORs have a unique signaling profile that shares little overlap with the responses initiated by ORs in the PM. OR activation at the Golgi apparatus by permeant drugs leads to minimal effects on gene transcription yet has a unique imprint on the phosphoproteome.

## Discussion

The present study establishes that activation of ORs in different cellular compartments promotes distinct signal transduction. Focusing on MOR and DOR in the PM and in the Golgi apparatus, we uncover similarities and differences in receptor-proximal protein recruitment, and delineate location-specific effects on gene expression and protein phosphorylation. The results convey that intracellular ORs do not merely mimic the actions of PM receptors, but that they uniquely drive cellular responses upon activation by permeant drugs. We find that OR coupling can be promoted by transducer–lipid interactions that depend on the local membrane composition, which suggests a key role for lipids in regulating GPCR coupling with subcellular precision.

To interrogate OR-proximal protein engagement in a location-resolved manner, we focused our analyses on mG probes, GRK2/3, and β-arrestins, which are cytosolic proteins and thus readily available to bind ORs in both locations. For the first time, we characterize and employ Gαi-based probes to study GPCR coupling, and detect that mGi1, mGi2, and mGi3 rapidly bind active ORs in PM and Golgi apparatus. While mGi interaction relies on agonist-driven GPCR conformational changes, β-arrestin binding additionally requires ORs to be phosphorylated (*31, 52*). We here discover that Golgi-localized MOR and DOR are phosphorylated by GRK2/3 in an agonist-dependent manner, however they do not drive β-arrestin recruitment, contrasting with PM ORs in the same cell. It conveys that additional, location-specific determinants regulate β-arrestin–OR binding (*32*). β-arrestins can directly interact with membrane lipids via loops in the C-domain, which contributes to the stability of select β-arrestin–GPCR complexes (*53, 54*). In particular, the PM phosphoinositide PIP2 is critical for β-arrestin binding to a subset of GPCRs, including MOR and DOR (*55*). Our data on OR–mGsi coupling suggests that lipid-based selection mechanisms may be widely relevant for GPCR–transducer coupling and provide a rationale for the absence of mGsi and β-arrestin2 engagement with MOR and DOR in the Golgi apparatus, which lacks PIP2.

The MD simulations of MOR–mGi1 and MOR-mGsi complexes embedded in membrane models with PM- or Golgi-like lipids reveal that mGsi, but not mGi1, binds MOR in a membrane-dependent manner. The findings show that two highly similar GPCR binding proteins can remarkably differ in their sensitivity to lipid regulation, which has broader implications for G protein coupling selectivity by GPCRs: mGsi is a chimeric probe that is almost entirely based on Gαs, which suggests that the lipid sensitivity may be a specific determinant of Gαs–GPCR engagement. Consistent with this, a recent study found reduced mGs recruitment to Golgi-localized dopamine D1 receptors when compared to the PM receptor pool (*56*). Furthermore, mass-spectrometry-based experiments identified PIP2 as a lipid that enhances mGs coupling to adenosine A2A receptors and found that mGs binding to PIP2 stabilized the interaction with β1 adrenergic receptors (*57*). It suggests that the rules for GPCR coupling selectivity may differ at intracellular organelles relative to the PM according to the lipid sensitivities of individual G proteins.

The present results, together with previous studies, establish that ORs in the Golgi apparatus are signaling competent as they engage transducer proteins (G proteins, GRKs) and inhibit cAMP production in response to permeant agonists (*7, 32*). Yet the transcriptomic and phosphoproteomic analyses delineate prominent differences in the downstream effects of Golgi-localized DOR relative to the PM receptor pool. A striking finding is that activation of DOR in the Golgi apparatus is uncoupled from transcriptional effects 1.5 h and 6 h after agonist activation, in stark contrast to PM OR signaling. Yet transcription factors account for 20% of phosphoproteins modulated by Golgi-localized OR signaling. We note that several of the identified transcription factors, including STAT3 and FOXC2, are known to require multisite phosphorylation for altered activity (*58–60*). Multisite regulation was, however, not detected in our phosphoproteomics dataset. Therefore, we speculate that subtle phospho-regulation of transcription factors by Golgi-localized DOR signaling may not propagate to the transcriptional level due to noise-filtering in the signal transduction cascade (*61, 62*). Golgi-localized OR signaling also led to phospho-regulation of a large proportion of RNA binding proteins (RBPs), which are involved in mRNA translation, decay, splicing, or export (*63*). It is therefore possible that OR signaling at the Golgi apparatus promotes unique perturbation in mRNA homeostasis, and highlights an avenue of potential significance for future study of opioid drug-specific effects.

In addition to lipid-regulated mechanisms, location-specific GPCR signaling may also be promoted by differences in transducers and effector availability. This applies to heterotrimeric G proteins, whose Gα and Gγ subunits are anchored to membranes by protein lipidation. Different Gα subunits are found at the Golgi apparatus, yet Gβγ subunits predominantly localize at the PM under basal conditions (*64–67*). A greater number of Gαi/o heterotrimers in the PM relative to the Golgi compartment likely impacts the G protein-mediated signal strength. Of note, non-canonical signaling via Gα can occur at the Golgi apparatus, when Gα subunits signal alone or in complex with other proteins, such as guanine exchange modulators, upon GPCR activation (*67–69*). An example is the KDEL receptor, an atypical GPCR, that signals at the Golgi apparatus via Gαo subunits in complex with Rab1/Rab3 GTPases and αGDI (*70*). The phosphoproteomic signature downstream of Golgi-localized DOR activation suggests that non-canonical coupling mechanisms may contribute to the local signaling response. Another family of important OR interacting proteins comprises the GPCR kinases (GRKs), of which four (GRK2, 3, 5, and 6) are ubiquitously expressed and subdivided into cytosolic (GRK2/3) and the PM-associated (GRK5/6) proteins (*71*). Our data on Golgi-localized MOR, DOR, and β2AR phosphorylation indicates that regulation by the cytosolic GRK2/3 can occur at both PM and Golgi apparatus. However, GPCRs that underlie regulation by PM-anchored GRK5/6 are unlikely to be regulated by the same kinases at internal organelles, raising the possibility of altered GPCR–kinase coupling that may further promote location-selective effects.

Our work delineates Golgi-localized OR signaling in response to permeant opioid drugs in a simplified cellular system and focuses on effects that are initiated upon OR activation. OR family members couple to inhibitory Gi/o proteins, reduce cAMP production, inhibit calcium conductance, and promote inhibition of neuronal activity in vivo (*72*). Therefore, future studies should address if Golgi-localized OR signaling downstream of opioid drugs, when paired with a separate priming stimulus, may lead to additional and synergistic signaling responses that were not captured here. This is important given the role of ORs and their ligands in the neuromodulation of dynamic, transient, and fluctuating neuronal signals.

In sum, and viewed more broadly, the present results reinforce an emerging understanding that GPCRs in intracellular organelles contribute to specific agonist actions and promote signal events that differ from the canonical response initiated at the cell surface. We propose from the present observations that lipid-based regulatory mechanisms contribute to location-biased OR–transducer selection and directly impact functional selectivity. Although the physiological consequence of Golgi-localized drug-specific OR signaling remains concealed for the time being, our results provide novel molecular insights that advance the notion that different subcellular receptor pools contribute to functional selectivity in opioid drug action.

## Material and Methods

### Mammalian cell culture conditions

HEK293 (CRL-1573, ATCC, female) and HeLa cells (CRM-CCL-2, ATCC, female) were cultured in Dulbecco’s modified Eagle’s medium (DMEM, GIBCO), supplemented with 10% fetal bovine serum (FBS, GIBCO). HEK293 cells stably expressing N-terminally FLAG-tagged DOR or MOR were cultured in the presence of 250 μg/ml Geneticin (Gibco). For transient DNA expression, Lipofectamine 2000 (Invitrogen) was used according to the manufacturer’s instructions, transfection media was exchanged after 6h.

### Confocal microscopy-based protein recruitment assay

HeLa cells were seeded on poly-L-lysine-coated 35 mm Cellvis glass-bottomed dishes (IBL, 220.110.022) and after 24 h transfected with fluorescently-tagged receptors (MOR-GFP or SEP-DOR, 1 μg DNA), Ruby2-mG probes (0.6 μg) or β-arrestin2-mCherry (0.8 μg), and ManII-BFP (0.25 μg) using 3 µL Lipofectamine 2000. Cells were live-imaged 16-24 h post transfection in HBS imaging solution (Hepes buffered saline with 135 mM NaCl, 5 mM KCl, 0.4 mM MgCl2,1.8 mM CaCl2, 20 mM Hepes, 1 mM d-glucose, 1% FBS, adjusted to pH 7.4). Recruitment of mG probes and β-arrestin2 was monitored with a Spinning disk confocal microscope (Nipkow, Zeiss) using an EC Plan Neofluar 100x/1.3 Oil Ph3 objective in a temperature and CO_2_-controlled environment (37°C, 5% CO2). Images of the same cells were acquired before and 5 min after agonist addition. Fluorescence intensities of mG probes and β-arrestin2 in the Golgi area were measured (see quantitative image analysis) and normalized to OR fluorescence. Protein recruitment to Golgi-localized ORs is represented as the agonist-induced fold change relative to before agonist addition.

### Split NanoLuc-based protein interaction assay

HeLa cells were seeded into 6-well plates and individual wells transfected with i) 0.4 μg DOR-SmBiT and 0.25 μg LgBiT-mG probe, or ii) MOR-SmBiT, 1 μg FLAG-MOR, and 0.25 μg LgBiT-mG probe or iii) 0.4 μg DOR-SmBiT, 0.25 μg β-arrestin-LgBiT, and 0.8 μg GRK2-GFP, using 3 µL Lipofectamine 2000. Binding of mG probes or β-arrestin2 to ORs triggers NanoLuc (SmBiT and LgBiT) complementation. 16-24 h after transfection, cells were seeded into black, clear-bottom 384-well plates (20,000 cells/well) in DMEM without phenol red, containing 30 mM HEPES (pH 7.4) and the NanoLuc substrate Nano-Glo (Promega, N2012), and incubated for 45 min at 37°C. The luminescence signal was recorded using the FDSS/μCELL kinetic plate imager (Hamamatsu) with an integrated simultaneous dispensing head and simultaneous detection across the plate. After acquiring baseline luminescence for 3 min, agonists (concentration specified in figure legends) were added to the cells. Luminescence was recorded every 2 s for 5 min post agonist addition. Experiments were performed in the absence or presence of 10 µM CTAP (MOR antagonist) or 100 µM ICI (DOR antagonist) to block PM ORs (addition 5 min prior to acquisition). Luminescence values were normalized to baseline signal (before agonist) and to vehicle treated control cells. Recruitment kinetics are represented as a percentage of the maximal luminescence signal detected in the absence of antagonists.

### Immunofluorescence-based GPCR phosphorylation assay

HeLa cells were seeded onto 15 mm^2^ glass coverslips in 12-well plates and after 24 h transfected with FLAG-tagged GPCRs (FLAG-MOR, FLAG-DOR, or FLAG-β2AR, 0.4 μg DNA) and ManII-BFP (0.15 μg DNA) using 1.5 µL Lipofectamine 2000. GRK2-GFP (0.5 μg DNA) was co-transfected in selected DOR and β2AR experiments as indicated in figure legend. For experiments with ICI addition, only high GRK2-GFP expression cells were selected for quantification. 16-24 h after transfection, cells were treated with ligands (details in figure legends) for 5 min and fixed using 4% formaldehyde in PBS. To assess reversibility of MOR phosphorylation, morphine was washed out after 5 min and 100 μM Naloxone was added for 15 min before fixation. To probe the role of GRK2/3, 30 μM Comp101 was added to cells 45 min prior to agonist treatment. When indicated, 100 μM of ICI was added 5 min prior to agonist treatment. Cells were permeabilized and blocked with 0.1% Saponin and 1.5% BSA in PBS and incubated overnight at 4°C with primary phospho-specific anti-GPCR antibodies and anti-FLAG antibodies (see key resources table for antibody references and dilutions) in blocking solution. After three washes, cells were incubated with Alexa Fluor-labeled secondary antibodies (Thermo Fisher, 1:1000 dilution) for 45 min at room temperature. Samples were imaged with a spinning disk confocal microscope (Nipkow, Zeiss) using an EC Plan Neofluar 100x/1.3 Oil Ph3 objective. Signal of GPCR phosphorylation in the ManII-labeled Golgi area is normalized to Golgi-localized OR signal and to non-treated control cells.

### TIR-FM based protein recruitment assay

HeLa cells were seeded on poly-L-lysine-coated 35 mm glass-bottomed dishes and transfected with FLAG-DOR or FLAG-MOR (1 μg DNA) and mG probes (0.6 to 0.8 μg DNA) using 3 µL Lipofectamine 2000. 16-24 h after transfection, surface ORs were labeled for 10 min with anti-FLAG M1-AF647 and media changed to HBS imaging solution. Cells were imaged at 37°C using a Nikon Eclipse Ti microscope using a 100x 1.49 Oil CFI Apochromat TIR-FM objective, temperature chamber, objective heater, perfect focus system and an ORCA-Fusion BT Digital CMOS camera. Images were acquired before and 5 min after agonist addition. Protein relocalization (ΔF) was calculated as F(t)/F_0_ with F(t) = mG signal after ligand addition and F_0_ = mG signal before ligand addition, both normalized to OR fluorescence.

### Quantitative fluorescence image analysis

Unprocessed images were analyzed using ImageJ (2.1.0). Phosphorylation of GPCRs in the Golgi apparatus was quantified using a custom-written macro. The ManII signal was used as the Golgi mask and within the mask, the mean fluorescence signal of i) phosphorylated GPCRs (pGPCR) and ii) total GPCRs was measured. A zone around the ManII mask was generated to measure the cytosolic background signal. We then calculated the ratio of background-subtracted p-GPCR/GPCR signals for all acquired cells. Quantification of mG recruitment to ORs in living cells imaged with spinning disk confocal microscopy before and after agonist addition was performed in a similar manner: ManII was used as Golgi mask and fluorescence of i) mG and ii) OR measured in all cells. A region outside the cells was used as the background and background-subtracted mG/OR ratios were calculated. The mG recruitment level F/F_0_ was determined by normalizing the mG/OR level after agonist addition with the level before agonist addition. β-arrestin-mCherry recruitment to ORs was measured as follows: Spinning disk confocal microscopy images with focus on the Golgi plane were used to quantify β-arrestin and DOR signals in the ManII-defined Golgi mask. β-arrestin/DOR and DOR signals were measured before (F_o_) and after (F) agonist addition and recruitment signal calculated as F/F_0_. Confocal images with a focus on the cell surface were used to quantify β-arrestin and DOR signals at the PM. A mask encompassing each cell was generated and F/F_0_ calculated similar to the Golgi recruitment. To quantify mG recruitment to the PM based on TIR-FM images, a mask encompassing each cell was generated. The fluorescence intensity ratio of background-subtracted mG/OR in this mask was calculated. A region outside the cell was used as the background and quantification was performed before (F_o_) and 5 min after (F) agonist addition.

### Statistics of image quantification

Quantification of data is presented as mean ± standard deviation (SD) based on at least 3 biologically independent experiments with the precise number indicated in the figure legends. Statistical analysis was performed using Prism (9.1.1) and using unpaired, paired two tailed Student’s t test or one-way ANOVA test as indicated.

### Sample preparation for RNA sequencing

HEK293 cells stably expressing FLAG-DOR were treated for 1.5 h or 6 h with i) vehicle control ii) 100 nM DPDPE or iii) 100 nM SNC80 or iv) 100 nM SNC80 after 5 min pre-incubation with 100 μM ICI 174,864 (ICI present during entire experiment duration), with N=3 for all samples. Total RNA was extracted using the RNeasy Mini Kit (Qiagen), genomic DNA was removed using the RNase-free DNase kit (Qiagen) and samples analyzed using a bioanalyzer. Gene expression in 1.5 h samples was measured using 3’ RNA sequencing (UC Davis Genome Centre) and in 6 h samples using whole transcript RNA sequencing (iGE3 Genomics Platform, University of Geneva). Multiplexed libraries for the 1.5 h samples were generated using the Lexogen QuantSeq 3’ mRNA-Seq Library Prep Kit and for the 6 h samples using TruSeqHT Stranded Total RNA Library Prep protocol (Illumina).

### RNA sequencing and analyses

Libraries generated for the 1.5 h treatment were sequenced using the Illumina NextSeq 500 or HiSeq 4000 devices with a single-end read length of 80-90 nucleotides yielding at least 3 millions mapped reads per sample. Sequences were mapped to the human genome GRC h38.12 using GENCODE v28 annotation with STAR v. 2.6.0c. Generated raw counts were normalized and genes with low expression were filtered out leaving 15,946 genes. Differential expression analyses and related statistics between control and treated conditions were conducted using the Limma-voom R package 3.38.2. Libraries generated for the 6 h treatment were sequenced using the Illumina HiSeq 4000 device with a single-end read length of 50 nucleotides yielding around 22 million mapped reads per sample. Fastq files were generated using Illumina bcl2fastq v. 2.20.0 and mapped to the human genome UCSC hg38 using STAR v. 2.7.0f. Aligned reads were quantified with the Python software htseq-count v. 0.9.1. Genes were filtered down to 14,467 after removal of low expressers and normalized using the R package EdgeR 3.28.1. Differential gene expression between control and treated conditions was computed and statistics were performed using the General Linear Model, quasi-likelihood F-test (QL F-test) with FDR and Benjamini & Hochberg correction. Any gene with fold-change over 1.5 or below −1.5 and p-value < 0.05 was considered as significantly regulated.

### Sample preparation for mass spectrometry

HEK293 cells stably expressing FLAG-DOR were treated for 5 min or 25 min with 100 nM SNC80 in the presence or absence of ICI (5 min pre-incubation and present throughout the experiment), with N=3 for all samples. Cells were then washed with ice-cold PBS, spun down, and dried pellets snap-frozen in N_2_. Thawed pellets were lysed in a buffer containing 50 mM Tris pH 7.4, 25 mM NaCl, 2% SDS, 2.5 mM EDTA, 20 mM TCEP and 2% SDS, supplemented with Halt-protease and Halt-phosphatase inhibitors (Thermofisher). Samples were vortexed and heated at 95°C for 10 min with 400 rpm mixing on a thermomixer. DNA was sheared by sonication. Samples were centrifuged for 30 min at 17’000 g, the supernatant was collected and incubated with 0.5 M iodoacetamide for 1 h at room temperature. Proteins were digested based on the FASP method (Wiśniewski et al. 2009) using Amicon Ultra-4, 30 kDa centrifugal filter units (Millipore). Trypsin (Promega) was added at 1:75 enzyme/protein ratio and digestion was performed overnight at room temperature. The resulting peptide samples were desalted with C18 macrospin columns (Harvard Apparatus) and then completely dried under speed-vacuum. Each sample was labeled with corresponding TMT-9plex reagent dissolved in 110 μl of 36% CH_3_CN, 200 mM EPPS (pH 8.5). Reaction was performed during 1 h at room temperature and quenched by adding hydroxylamine to a final concentration of 0.3% (v/v). Labeled samples were pooled, dried and desalted with a peptide desalting spin column (Thermo Fisher). Phosphopeptides were enriched using the High-Select Fe-NTA Phosphopeptide Enrichment Kit (Thermo Fisher). Phosphopeptide and the flow-through fractions were desalted with a C18 macrospin column (Harvard Apparatus) and completely dried under speed-vacuum. The flow through fraction was then fractionated into 13 fractions using the Pierce High pH Reversed-Phase Peptide Fractionation Kit (Thermo Fisher).

### Quantitative real-time PCR

HEK293 cells stably expressing FLAG-DOR were treated with 100 nM DOR agonist DADLE for 1.5 h. Total RNA was extracted and genomic DNA was removed as described above. RNA samples were quantified using the Nanodrop device (Thermo Fisher) and subjected to reverse transcription using the High-Capacity cDNA Reverse Transcription Kit (Applied Biosystems). RT-qPCR was performed using the PowerUp Sybr green reagent (Applied Biosystems) and relative RNA quantification was performed using the Delta-Delta Ct method.

### ERK1/2 activation assay

HEK293 cells stably expressing FLAG-DOR were treated with 100 nM SNC80 in the presence or absence of ICI 174,864 (5 min pre-incubation and present throughout experiment) for the indicated times. Cells were lysed in RIPA buffer (10 mM Tris pH 7.4, 1X Triton, 150 mM NaCl, 0.1% SDS, 2 mM EDTA) supplemented with protease and phosphatase inhibitors (Roche) and lysates were sonicated followed by quantification using the Pierce BCA protein assay (Thermo Fisher). Protein samples were mixed with NuPAGE LDS sample buffer and 100 mM DTT, and heated for 10 min at 70°C. Lysates were separated by SDS-PAGE using 4-12% Bis-Tris Plus gels (Thermo Fisher) and transferred to an isopropanol-activated PVDF membrane. After blocking in TBS with 5% BSA, the membranes were incubated overnight at 4°C with anti-ERK1/2 and anti-phosphorylated-ERK1/2 primary antibodies (1:2000) in TBS-Tween (0.1% v/v) with 5% BSA. After washing, HRP-coupled anti-mouse and anti-rabbit secondary antibodies were added for 1 h at RT. Membranes were imaged in the presence of Pierce ECL SuperSignal West Pico plus (Thermo Fisher) using the iBright 1500 device (Invitrogen). ERK protein levels were quantified using ImageJ and pERK/ERK was plotted.

### Mass spectrometry and analyses

Phosphopeptides were reconstituted in loading buffer (5% CH_3_CN, 0.1% FA) and 1 μg was injected into the column. LC-ESI-MS/MS was performed on an Orbitrap Fusion Lumos Tribrid mass spectrometer (Thermo Fisher) equipped with an Easy nLC1200 liquid chromatography system (Thermo Fisher) at the Proteomics Core Facility, University of Geneva. Peptides were trapped on a Acclaim pepmap100, C18, 3μm, 75 μm x 20 mm nano trap-column (Thermo Fisher) and separated on a 75 μm x 500 mm, C18 ReproSil-Pur from Dr. Maisch GmBH, 1.9 μm, 100 Å, home-made column. The analytical separation was run for 125 min using a gradient of H_2_O/FA 99.9%/0.1% (solvent A) and CH_3_CN/FA 80%/0.1% (solvent B). Data-Dependent Acquisition (DDA) was performed with MS1 full scan at a resolution of 120’000 FWHM followed by as many subsequent MS2 scans on selected precursors as possible within 3 second maximum cycle time. High pH Reversed-Phase Peptide fractions were treated similarly. Raw data were processed using Proteome Discoverer (PD) 2.3 software (Thermo Fisher). In brief, spectra were extracted and searched against the Human Reference Proteome database (release 11_2019, 20’660 entries) combined with an in-house database of common contaminants using Mascot (Matrix Science, London, UK; version 2.5.1). Trypsin was selected as the enzyme, with one potential missed cleavage. Precursor ion tolerance was set to 10 ppm and fragment ion tolerance to 0.02 Da. Carbamidomethylation of cysteine (+57.021) as well as TMT10plex (+229.163) on lysine residues and on peptide N-termini were specified as fixed modification. Oxidation of methionine (+15.995) as well as phosphorylated (+79.966) serine, threonine, and tyrosine were set as variable modifications. Search results were validated with a Target Decoy PSM validator. PSM and peptides were filtered with a false discovery rate (FDR) of 1%, and then grouped to proteins with again a FDR of 1% and using only peptides with high confidence level. Both unique and razor peptides were used for quantitation and protein and peptide abundances values were based on S/N values of reporter ions. The abundances were normalized on “Total Peptide Amount” and then scaled with “On all Average”. All the protein ratios were calculated from the medians of the summed abundances of replicate groups and associated p-values were calculated with an ANOVA test based on individual proteins or peptides. Any phosphopeptide with a fold-change ratio over 1.2 or below −1.2 and p-value < 0.05 was considered as significantly regulated.

### Phosphoproteomics analyses and upstream kinase predictions

Upstream kinase predictions were performed using the KSEA app (v. 1.0, (*73*)) fed with a pooled list of unique phosphopeptides significantly regulated at 5 min and 25 min upon SNC80 or ICI-SNC80 treatment. Kinase plots were generated with a NetworKIN score cutoff set to 1, a substrate count cutoff set to 5, and a p-value cutoff set to 0.05.

### MD simulations

The 3D structures of the MOR (Uniprot ID: P35372) complexed with mGi1 and mGsi were built using “*Alphafold2-multimer*” (*74*). Subsequently, the two heterodimers were embedded into tailored phospholipid bilayers using CHARMM-GUI (*75, 76*) in order to obtain the following four systems: 1) MOR-mGi1 in PM, 2) MOR-mGsi in PM; 3) MOR-mGi1 in Golgi-like membrane (GM); 4) MOR-mGsi in GM. To reproduce the properties of PM and GM (*36–38*), two different phospholipidic compositions were employed, which are detailed in Figure 5A. All systems were solvated using a box of TIP3P water model and a salinity of 0.15 M NaCl. The N-terminus and C-terminus of MOR, mGi1, mGsi were capped with an acetyl and a metil-amino protecting group, respectively. The CHARMM36m force field was employed for the MD simulations (*77*), which were run using the GROMACS 2021.5 engine (*78*). Each simulation box underwent a thermalization cycle with smoothly decreasing restraints on heavy atoms to gently equilibrate the structure. We employed the following protocol: heating the system from 100 K to 300 K by increasing the temperature by 50 K each step composed of 1 ns of NVT simulation followed by 1 ns of NPT simulation. During the thermalization, the V-rescale thermostat was employed, whereas the Langevin dynamics temperature control scheme was used in the production runs. Periodic boundary conditions were applied, and the particle-mesh-Ewald (PME) method was used to treat long-range electrostatic interaction (*79*). For short-range interactions, a cut-off distance of 1.0 nm was applied. The pressure was fixed at a reference value equal to 1 bar by means of the Parrinello-Rahman barostat (*80*).

### Cluster analysis

Cluster analyses on the MD trajectories were performed using GROMACS’s “*gmx cluster*” routine. The cluster families of MOR-mGi1 and MOR-mGsi were obtained by aligning on secondary structure’s Cα atoms of MOR and computing the RMSD for the secondary structure’s Cα of both mGi1 and mGsi. The RMSD cut-off was set at 1.5 Å.

### Binding interface evaluation

The protein-protein interactions established by the residues at the MOR-mGi1 and MOR-mGsi interface were assessed by calculating their frequency of occurrence using the “*PLOT NA*” routine of “*Drug Discovery Tool*” (DDT) (*81*) and displayed as histograms. We set a distance cut-off value of 4.0 Å to define two interacting residues.

### Table of key reagents and resources

| REAGENT or RESOURCE | SOURCE | IDENTIFIER |
| --- | --- | --- |
| **Antibodies** |  |  |
| Mouse anti-FLAG (M1) (1:1000) | Sigma-Aldrich | Cat# F-3040 |
| Phospho mu Opioid Receptor (Ser375) Primary Antibody (1:500) | Cell Signaling Technology | Cat# 3451S |
| Phospho delta Opioid Receptor (Ser363) Primary Antibody (1:1000) | 7TMantibodies | Cat# 7TM0317B |
| Phospho β2-Adrenergic Receptor (Thr360/Ser364) primary Antibody (1:250) | 7TM Antibodies | Cat# 7TM0029B |
| Human anti-Giantin (1:100) | Geneva antibody facility | ABCD-AA341 |
| Mouse anti-ERK1/2 (1:2000) | CST | 4696S |
| Rabbit anti-pERK1/2 (T202/Y204) (1:2000) | CST | 4370S |
| Mouse IG2A Anti-alpha Tubulin antibody (1:5000) | Geneva antibody facility | ABCD_AA345 |
| Goat anti-Rabbit IgG Secondary Antibody, Alexa Fluor 647 (1:1000) | Thermo Fisher Scientific | Cat# A32733 |
| Goat anti-Mouse IgG Secondary Antibody, Alexa Fluor 568 (1:1000) | Thermo Fisher Scientific | Cat# A11031 |
| Goat anti-Human IgG Secondary Antibody, Alexa Fluor 647 (1:1000) | Thermo Fisher Scientific | Cat# A21445 |
| HRP-anti-mouse (1:10000) | Sigma-Aldrich | A5278 |
| HRP-anti-rabbit (1:4000) | Sigma-Aldrich | A8275 |
| **Chemicals, Peptides, and Recombinant Proteins** |  |  |
| DAMGO, [D-Ala2, N-Me-Phe4, Gly5-ol]-Enkephalin acetate salt | Sigma-Aldrich | Cat# E7384 |
| DADLE, [D-Ala2, N-Me-Phe4, Gly5-ol]-Enkephalin acetate salt | Sigma-Aldrich | Cat# E7131 |
| DPDPE, [D-Pen<sup>2,5</sup>]-Enkephalin hydrate | Sigma-Aldrich | Cat# E3888 |
| Morphine sulfate | CAS #6211-15-0 |  |
| Fentanyl | CAS #1443-54-5 |  |
| SNC80 | Tocris | Cat# 0764 |
| ICI 174,864 | Tocris | Cat# 0820 |
| CTAP | Tocris | Cat# 1560 |
| Alexa Fluor 647 Protein Labeling Kit | Thermo Fisher Scientific | Cat# A20173 |
| ProLong Glass Antifade Mountant | Invitrogen | Cat# P36982 |
| Lipofectamine 2000 | Invitrogen | 11668019 |
| RNeasy Mini Kit | Qiagen | 74104 |
| RNase-Free DNase Set | Qiagen | 79254 |
| High-Capacity cDNA Reverse Transcription Kit | Thermo Fisher Scientific | 4368814 |
| PowerUp SYBR Green Master Mix | Applied Biosystems | A25741 |
| **Experimental Models: Cell Lines** |  |  |
| Human HEK293 cells | ATCC | CRL-1573 |
| Human HeLa cells | ATCC | CRM-CCL-2 |
| **Recombinant DNA** |  |  |
| pCAGGS/SE: ssf-MOR-EGFP, murine | (7) | N/A |
| pcDNA3.1: MOR-smBIT, murine | This study | N/A |
| pCAGGS-SE: SEP-DOR, murine | (82) | N/A |
| pcDNA3.1: DOR-smBIT, murine | This study | N/A |
| pcDNA3: signal sequence FLAG (ssf)-MOR, murine | Mark von Zastrow lab | N/A |
| pcDNA3: signal sequence FLAG (ssf)-DOR, murine | Mark von Zastrow lab | N/A |
| pcDNA3: signal sequence FLAG (ssf)-β2AR, murine | Mark von Zastrow lab | N/A |
| pCAGGS-SE: β-Arrestin2-mCherry, bos taurus | (83) | N/A |
| pNBe3: LgBIT-β-Arrestin2, human | (27) | N/A |
| pCAGGS-SE: GRK2-GFP, bos taurus | (82) | N/A |
| ManII-BFP2 | (70) | N/A |
| mRuby2-miniGo | Nevin Lambert | N/A |
| mRuby2-miniGsi | Nevin Lambert | N/A |
| mRuby2-miniGi1 | This study | N/A |
| mRuby2-miniGi2 | This study | N/A |
| mRuby2-miniGi3 | This study | N/A |
| LgBIT-miniGi1 | This study | N/A |
| **Software and Algorithms** |  |  |
| Prism | GraphPad | 9.1.1 |
| ImageJ | Imagej.net/contributors | 2.0.0-rc-54/1.51g |
| Excel | Microsoft | 16.0.10390.20024 |
| Illustrator CC | Adobe | 21.0.2 |
| Proteome Discoverer | Thermo Scientific | 2.4 |
| BioRender | biorender.com |  |

## Acknowledgements

We thank members of the Bioimaging Core Facility, the RE.A.D.S. Unit, the Proteomics Core Facility and the iGE3 Genomics Platform at the University of Geneva, and the Genome Centre at UC Davis for valuable discussion. We thank Nevin Lambert for sharing the mGo-mRuby2 construct. We thank Gonzalo Solis, Kevin Assoumou, Anne-Claude Gavin, Patrick Meraldi, and Simon Braun for valuable comments on the project and manuscript. We thank all group members for discussions and critical reading of the manuscript. The study was supported by the Swiss National Science Foundation Eccellenza Professorial Fellowship to MS (project number PCEFP3_181282) and a grant from the Olga Mayenfish Stiftung. We acknowledge PRACE and the Swiss National Supercomputing Centre (CSCS) for generous supercomputer time allocations on Piz Daint (project IDs: pr126, s1169). F.L.G. acknowledges the Swiss National Science Foundation and the BRIDGE program for financial support (project numbers 204795 and 203628). We thank Mark von Zastrow for discussions and supporting a part of this work at an early stage through NIDA NIH DA10711 and DA012864.

---

**Supplementary Figure S1:**
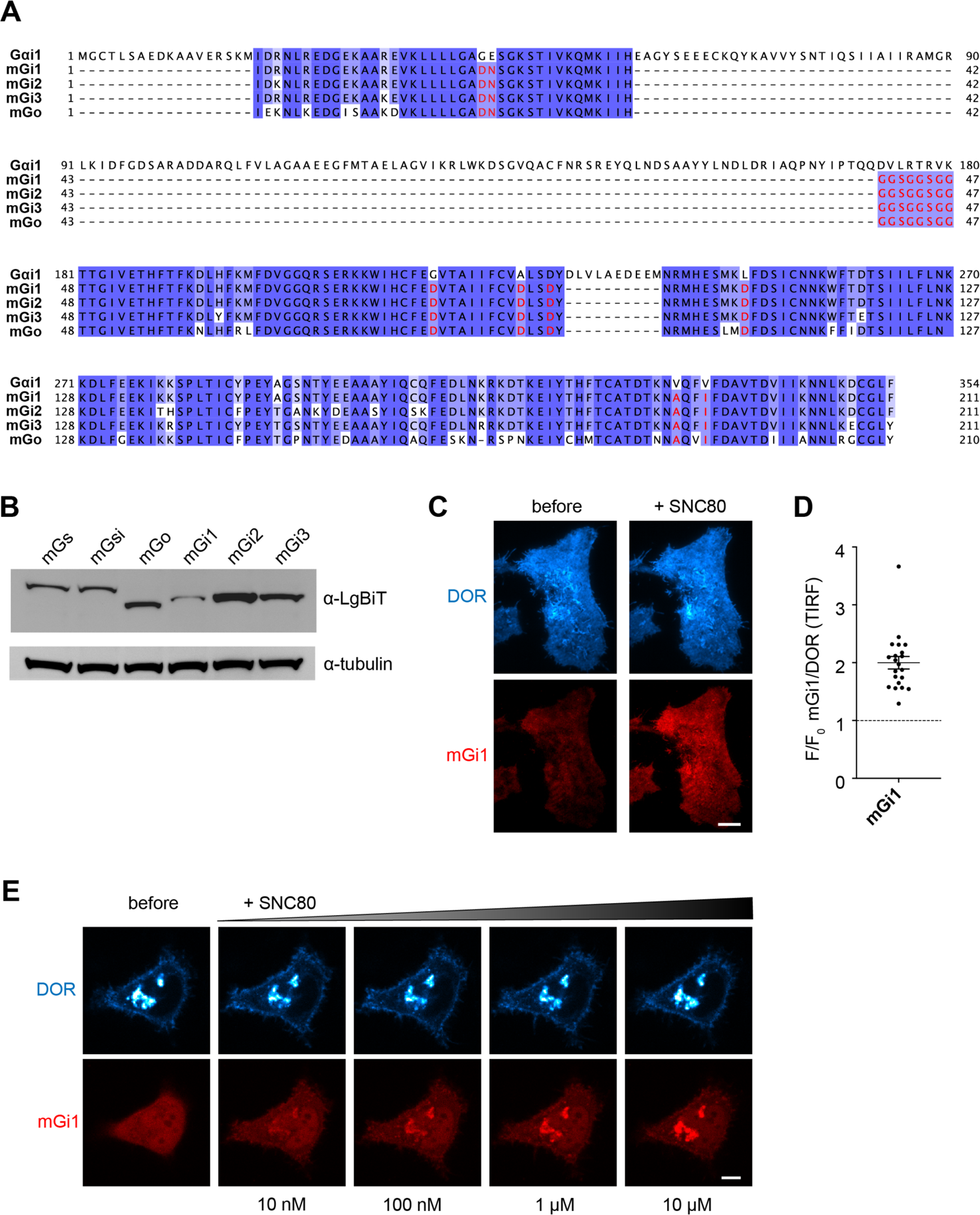
mG proteins derived from Gαi/o subunits serve as sensors for OR activation. **(A)** Protein sequence alignment of full-length human Gαi with mGi1, mGi2, mGi3 and mGo. Red amino acids indicate stabilizing mutations in the mGi/o proteins. **(B)** mG (mGs, mGsi, mGo, mGi1, mGi2, mGi3) protein levels quantified by Western blot. α-tubulin was used as a loading control. **(C)** TIR-FM images of living HeLa cells expressing FLAG-DOR (cyan, surface labeled with anti-FLAG M1-AF647) and mGi1-mRuby2 (red), before and 5 min after adding 10 µM SNC80. Scale bar = 10 μm. **(D)** Quantification of mGi1 intensity in TIR-FM images acquired before and 5 min after SNC80 addition, normalized to PM DOR signal. N=3 with > 20 cells analyzed, mean +/- SD. **(E)** Confocal images of living HeLa cells, expressing DOR-SEP (cyan) and mGi1-mRuby2 (red) before and after adding increasing concentrations of SNC80 (10 nM, 100 nM, 1 µM, 10 µM). Scale bar = 10 μm.

**Supplementary Figure S2:**
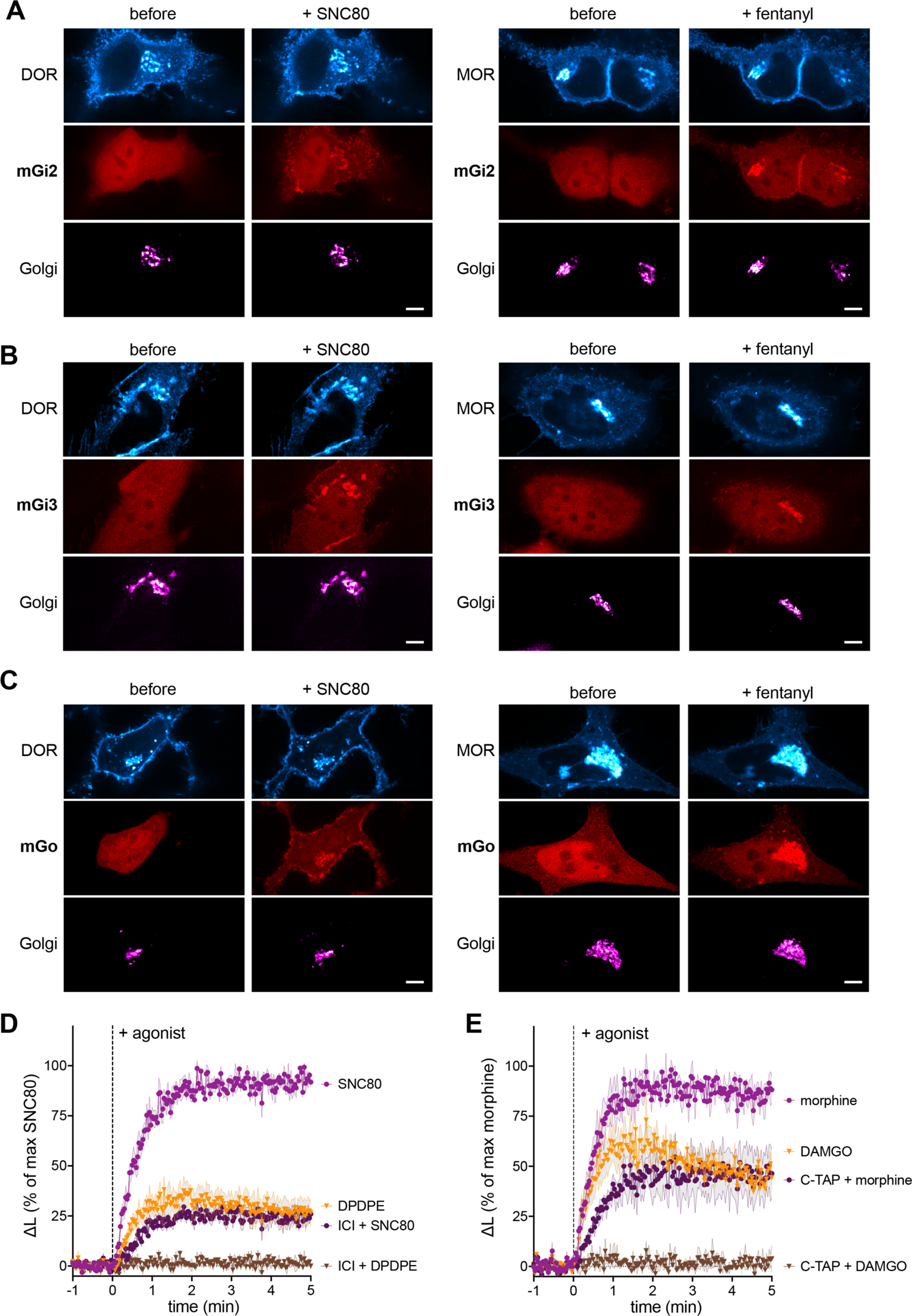
Activated Golgi-localized ORs recruit Gαi/o sensors. **(A)**, **(B)**, **(C)** Confocal images of living HeLa cells, expressing DOR-SEP (left panels, cyan) or MOR-GFP (right panels, cyan), mG proteins tagged with mRuby2 (red), and ManII-BFP (magenta) before and 5 min after 10 µM SNC80 or 1 μM fentanyl addition. Scale bar = 10 μm. **(A)** mGi2-mRuby2, **(B)** mGi3-mRuby2, **(C)** mGo-mRuby2 expression. **(D**) Change in luminescence signal through agonist-induced interaction of mGi1-LgBiT with DOR-SmBiT (left panel) or MOR-SmBiT (right panel). Data normalized to maximum signal upon SNC80 (left) or morphine (right) addition. Left: 100 nM DPDPE or SNC80 with or without pre-incubation with ICI (100 µM). Right: 100 nM DAMGO or morphine with or without pre-incubation with CTAP (10 µM). N=3, mean ± SEM.

**Supplementary Figure S3:**
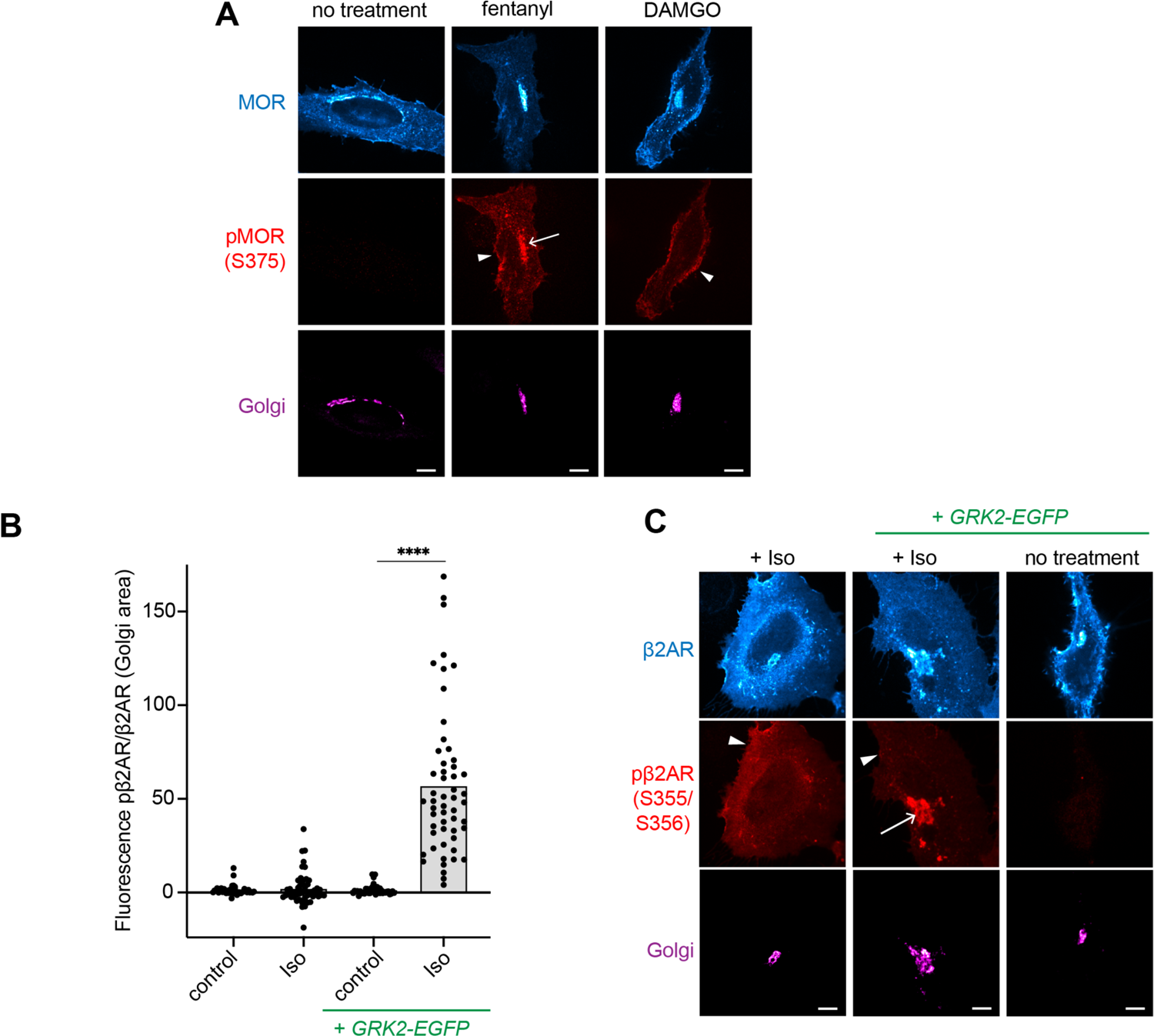
Activated GPCRs in the Golgi apparatus are regulated by phosphorylation. **(A)** Confocal images of HeLa cells expressing FLAG-MOR and ManII-BFP (magenta). Cells were fixed, permeabilized and immunolabeled with anti-FLAG (cyan) and anti-pMOR-S375 (red) antibodies. Cells were untreated, or treated with DAMGO (10 μM) or fentanyl (1 μM) for 5 min (quantification in Figure 2C). Scale bar = 10 μm. Arrow depicts pDOR in the Golgi area, arrowheads depict pDOR at the PM. **(B)** Quantification of pβ2AR/β2AR fluorescence in Golgi area (ManII-labelled). Cells were transfected and stained as in (C). Data normalized to signal of untreated control cells. Cells were treated with isoproterenol (Iso) for 5 min. Conditions with GRK2-EGFP co-expression are indicated. N=3 with cells analyzed >45. ****p<0.0001 by ordinary one-way ANOVA. **(C)** Confocal images of HeLa cells expressing FLAG-β2AR and ManII-BFP (magenta). Cells were immunolabeled with anti-FLAG (cyan) and anti-pβ2AR-S355/S356 (red) antibodies. Treatment with 10 μM Iso for 5 min of cells with or without GRK2-GFP expression. Arrow depicts pβ2AR in the Golgi area, arrowheads depict pβ2AR at the PM. Scale bar = 10 μm.

**Supplementary Figure S4:**
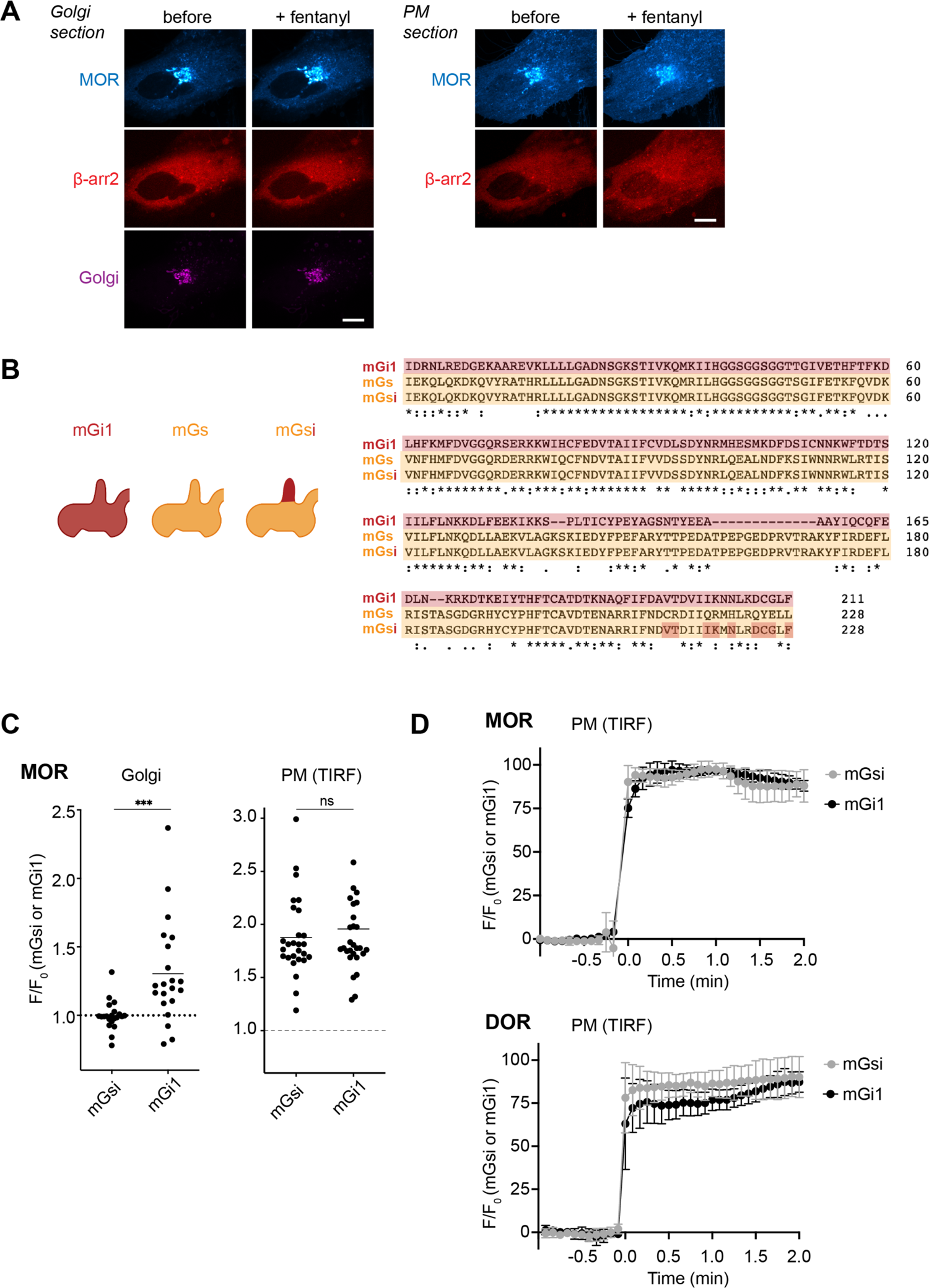
Location-selective OR binding to β-arrestin2 and mGsi. **(A)** Confocal images of living Hela cells, expressing GFP-MOR (cyan), β-arrestin2-mCherry (red), and a ManII-BFP (magenta) before and 5 min after 1 µM fentanyl addition. 2 confocal sections of the same cells are shown. Left: focus on the Golgi area, right: focus on the PM. Scale bar = 10 μm. **(B)** Left: Schematic showing differences and similarity between mGi1, mGs, and mGsi. Right: Protein sequence alignment of mGi1, mGsi, and mGs. **(C)** Quantification of mGsi-mRuby2 and mGi1-mRuby2 recruitment to MOR in Golgi or PM. Left: Probe recruitment to MOR in Golgi area (ManII-labeled) as measured by confocal imaging (normalized to Golgi MOR signal). Right: Probe recruitment to the PM as measured with TIR-FM imaging (normalized to MOR signal at PM). Same cells were imaged before and 5 min after 1 µM fentanyl addition. N=3 with > 15 cells analyzed, mean +/- SD. ***p<0.001 by unpaired t test. **(D)** Kinetic of mGsi (gray) and mGi1 (black) recruitment to ORs in the PM (top: MOR bottom: DOR) measured with TIR-FM. 5 s between frames is shown. F0, average fluorescence intensity before agonist. N=3, mean +/- SEM.

**Supplementary Figure S5:**
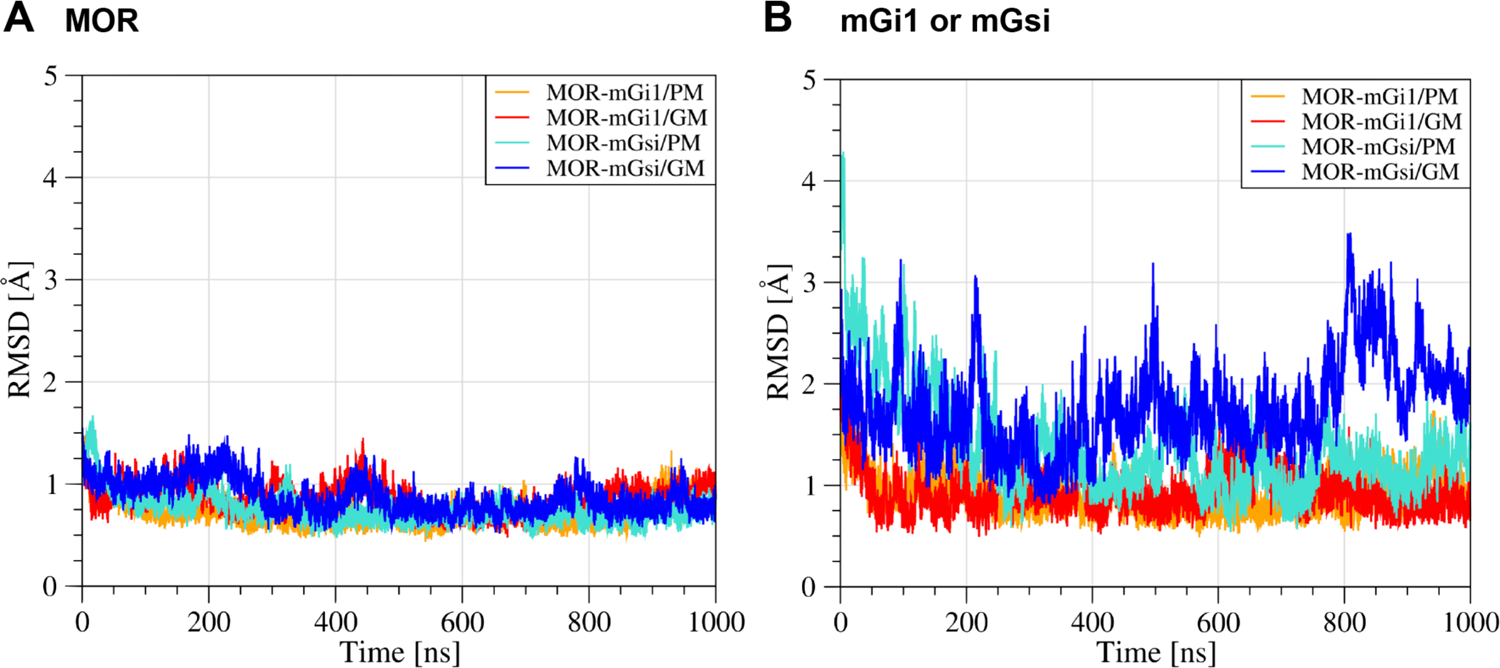
Dynamics and conformational stability of MOR, mGi1, and mGsi. RMSD plots extrapolated from the MD simulations performed on the MOR–mGi1 and MOR– mGsi heterodimers in PM and Golgi membranes. **(A)** Time evolution of MOR’s RMSD, measured on the secondary structure Cαs with respect to the average structure. **(B)** Time evolution of mGi1’s and mGsi’s RMSD, measured on their secondary structure Cαs with respect to each average structure.

**Supplementary Figure S6:**
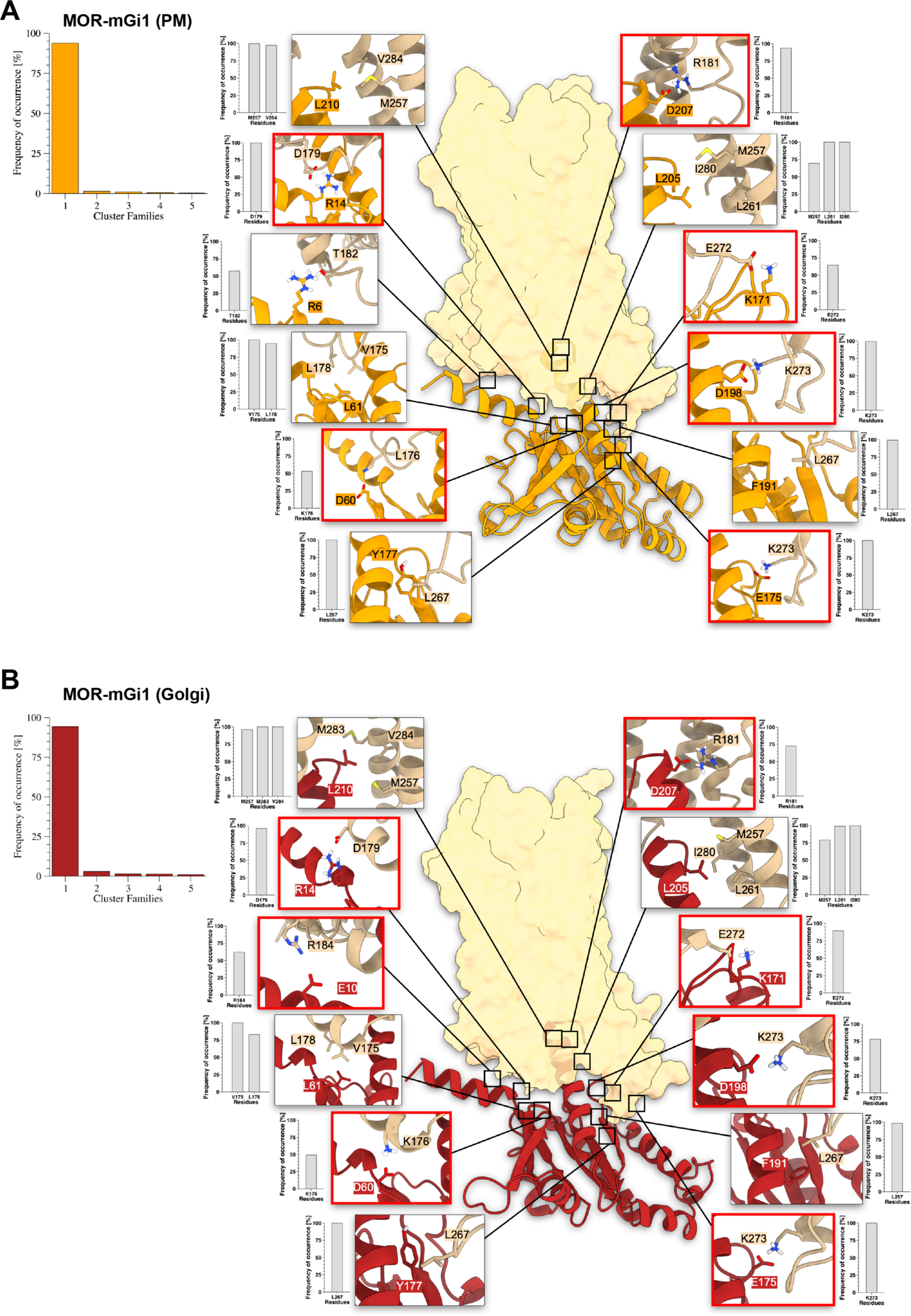
Structure analysis of PM-embedded and Golgi membrane-embedded MOR–mGi1 complexes. **(A)** The PM-embedded MOR–mGi1 heterodimer. Left: histogram of the conformational cluster analysis performed on the MD simulation. Right: MOR–mGi1 complex structure of the most populated cluster family. Insets show most relevant interactions between MOR and mGi1 and frequency of occurrence calculated from the MD simulation. Salt bridges highlighted with red box. **(B)** The Golgi membrane-embedded MOR–mGi1 heterodimer. Left: histogram of the conformational cluster analysis performed on the MD simulation. Right: MOR–mGi1 complex structure of the most populated cluster family. Insets show most relevant interactions between MOR and mGi1 and frequency of occurrence calculated from the MD simulation. Salt bridges highlighted with red box. MOR is colored in transparent yellow and represented through its solvent exposed surface. mGi1 in the PM or the Golgi lipid environment is colored in orange and red, respectively.

**Supplementary Figure S7:**
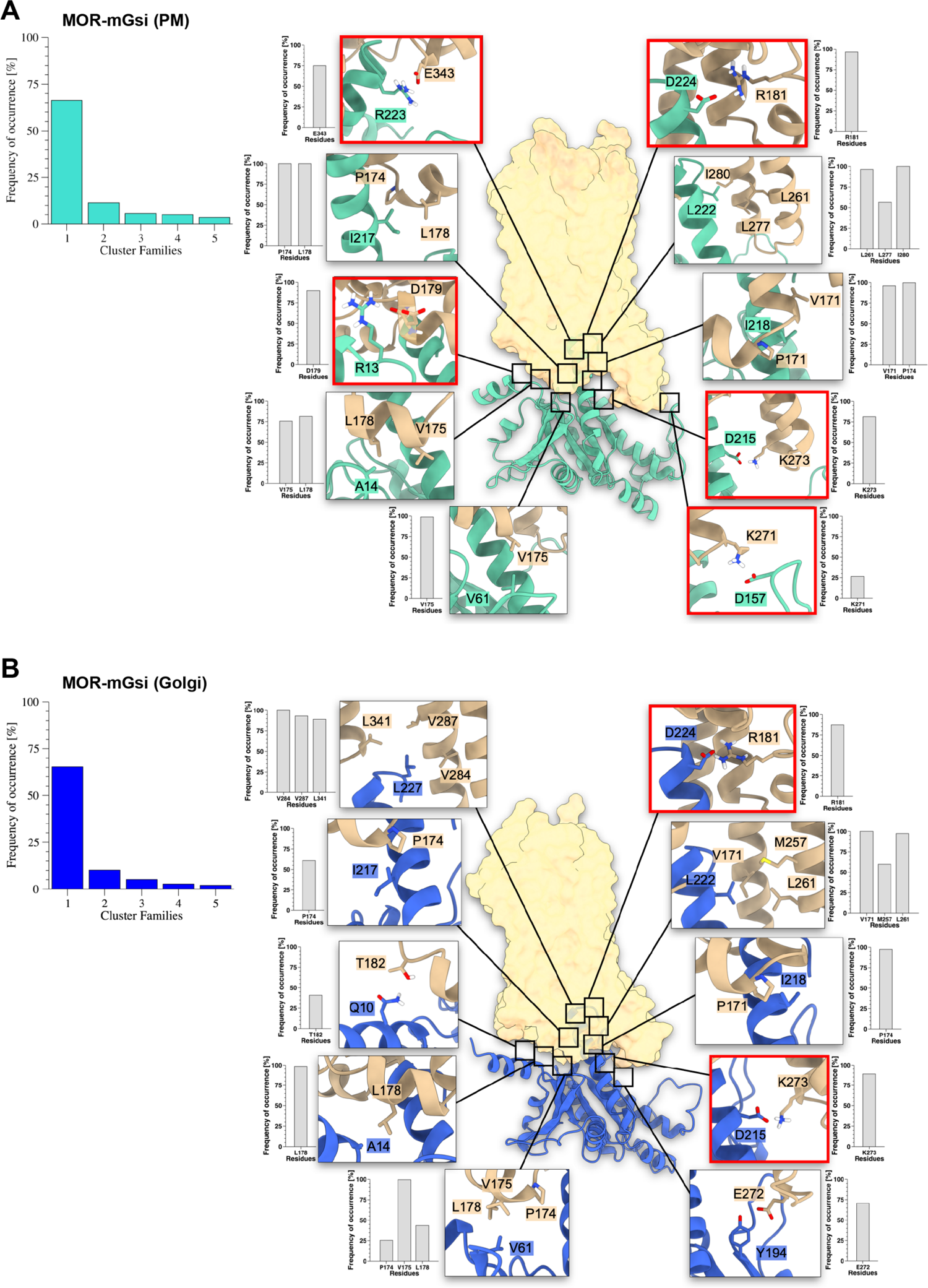
Structure analysis of PM-embedded and Golgi membrane-embedded MOR–mGsi complexes. **(A)** The PM-embedded MOR–mGsi heterodimer. Left: histogram of the conformational cluster analysis performed on the MD simulation; Right: MOR–mGsi complex structure of the most populated cluster family. Insets show most relevant interactions between MOR and mGsi and frequency of occurrence calculated from the MD simulation. Salt bridges highlighted with red box. **(B)** The Golgi membrane-embedded MOR–mGsi heterodimer. Left: histogram of the conformational cluster analysis performed on the MD simulation; Right: MOR–mGsi complex structure of the most populated cluster family. Insets show most relevant interactions between MOR and mGsi and frequency of occurrence calculated from the MD simulation. Salt bridges highlighted with red box. MOR is colored in transparent yellow and represented through its solvent exposed surface. mGsi in the PM or the Golgi lipid environment is colored in teal and blue, respectively.

**Supplementary Figure S8:**
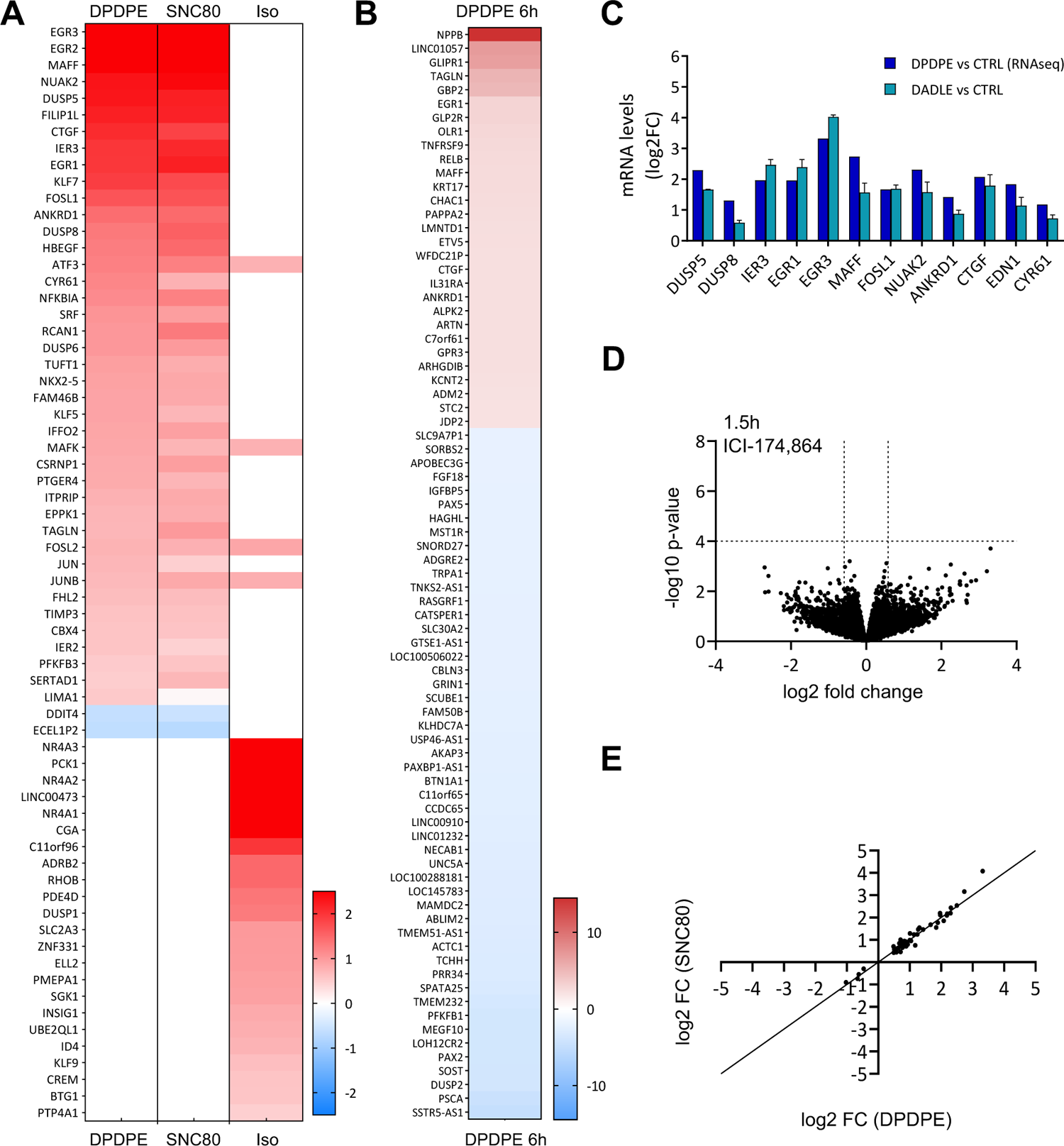
Characteristics of the DOR-driven transcriptional response. **(A)** Heatmap of differentially expressed genes in HEK293-DOR cells treated with DPDPE (100 nM) or SNC80 (100 nM) for 1. 5 h and upon β2AR activation with isoproterenol (Iso, 100 nM) for 1.5 h. Upregulated genes are shown in red (adj. p-value < 0.05) and downregulated genes are shown in blue (adj. p-value < 0.05). **(B)** Heatmap of differentially expressed genes in HEK293-DOR cells treated with DPDPE (100 nM) for 6 h. Upregulated genes are shown in red (FC > 2, adj. p-value < 0.05) and downregulated genes are shown in blue (FC < −2, adj. p-value < 0.05). **(C)** mRNA levels by RT-qPCR of selected genes regulated in HEK293-DOR cells upon treatment with DADLE (100 nM) for 1.5 h, compared to RNA-seq data. Relative gene expression was determined using the delta delta Ct method. GAPDH was used as reference gene. Results are presented as log2 fold-change over mock-treated control cells. N=2. **(D)** Volcano plots of differentially expressed genes in HEK293-DOR cells treated with ICI (100 µM) for 1.5 h versus mock-treated control cells. Mean differential gene expression from 3 replicates. Results are expressed as log2 fold-change. No genes are significantly changed in their expression (criteria as in Figure 6). **(E)** Scatter plot comparing the expression levels (log2FC) of regulated genes in HEK293-DOR cells after 1.5 h treatment with DPDPE (100 nM) or SNC80 (100 nM). The black line depicts x=y.

**Supplementary Figure S9:**
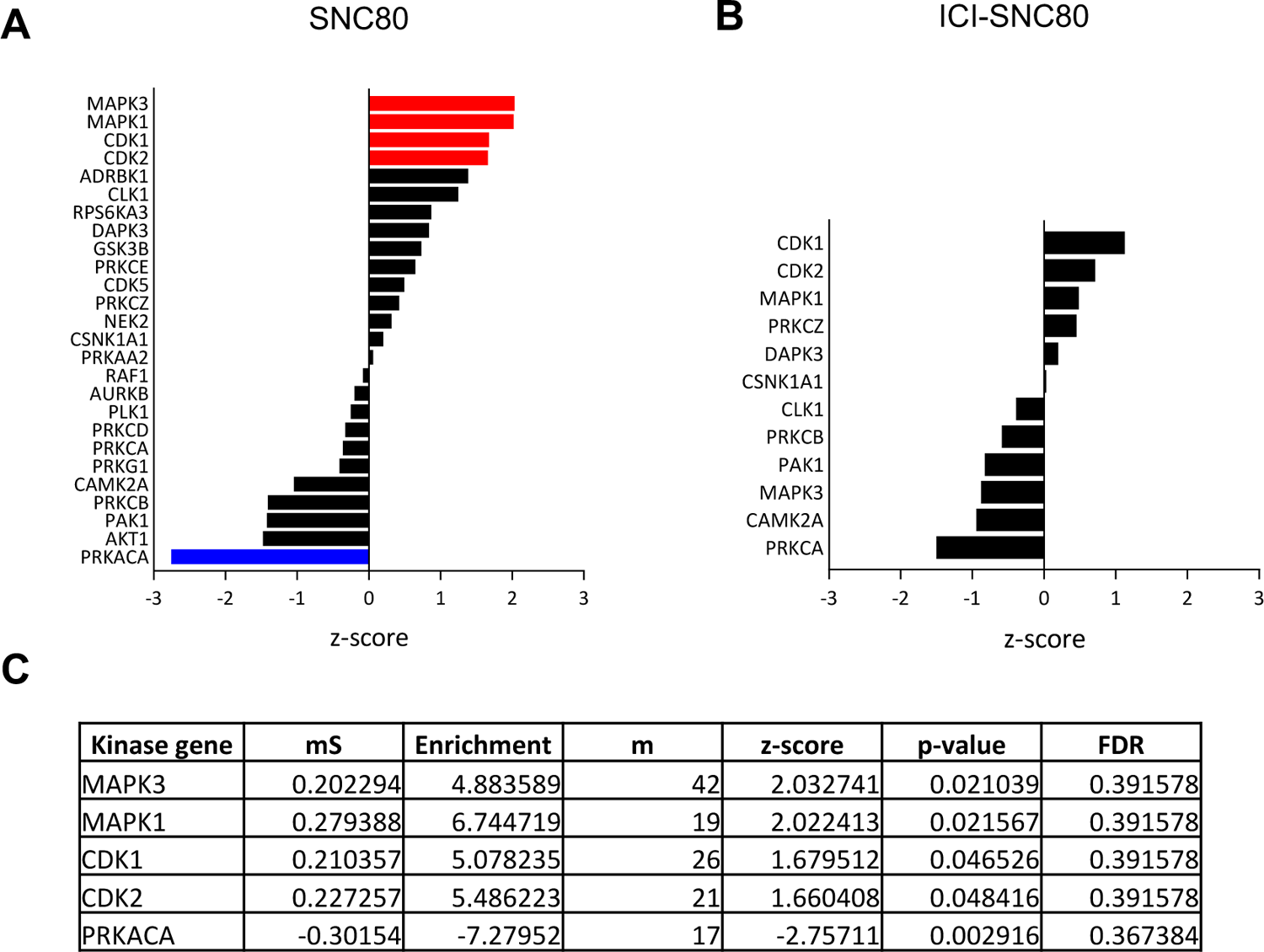
Kinase-substrate enrichment analyses for regulated phosphosites. **(A), (B)** Bar plots showing kinase activity predictions in HEK293-DOR cells treated with SNC80 **(A)** or ICI-SNC80 **(B)** as in Fig. 6A-D using significantly regulated phosphopeptides at both time points for each treatment condition. Results were generated using the KSEA app (PhosphoSitePlus + NetworKIN, NetworKIN score cutoff = 1, p-value cutoff = 0.05, substrate count cutoff = 5) and are presented as kinase z-score. Kinases predicted to have an increased activity compared to control are shown in red, kinases predicted to have a decreased activity are shown in blue. **(C)** Table summarizing the output data (kinases with p-value < 0.05) from the KSEA kinase prediction analysis upon SNC80 treatment presented in (A) (Kinase gene = kinase gene name, mS = mean Log2FC of all the kinase substrates, Enrichment = background-adjusted value of the kinase mS, m = total amount substrates detected from the experimental dataset for each kinase, z-score = normalized score for each kinase, FDR = adjusted p-value with Benjamini-Hochberg correction).

## Supplementary Tables

**Supplementary Table 1** (Excel): Transcriptomics data

**Supplementary Table 2 (Excel):** Phosphoproteomics data and analyses

